# A simple model captures key characteristics of biological non-deterministic genotype-phenotype maps

**DOI:** 10.1101/2025.07.23.666293

**Authors:** Nora S. Martin

## Abstract

By connecting genotypic mutations to the higher-order phenotypes relevant for selection, genotype-phenotype (GP) maps play a key role in evolution. GP maps are typically investigated using computational models of molecular phenotypes (for example, RNA secondary structures and simplified models of protein tertiary and quaternary structures), but GP map concepts are relevant beyond these specific models and so a systematic, model-independent approach is needed. This can be achieved by characterising GP maps in terms of properties like phenotypic frequencies, mutational robustness and evolvabilities, which can be computed for any given GP map. This approach has given insight into the shared features of GP maps and their evolutionary relevance. However, this progress is largely limited to the simplest case, where each genotype corresponds to a single, categorical phenotype. Here, I turn to a more realistic, but also more complex non-deterministic (ND) treatment, where each genotype generates an ensemble of phenotypes: I start by comparing the ND GP maps of the three biophysical models mentioned above using a recently proposed framework. Then, I find a simpler, synthetic map, which replicates key shared features for a range of modelling choices, suggesting that few ingredients are needed for these features to appear and highlighting the importance of non-linearities in GP maps. This synthetic ND GP map can be useful as a realistic, conceptually and computationally simpler model for future analyses of ND GP maps, their properties and their implications for evolutionary processes.

## I. INTRODUCTION

Variation through random mutations is a central ingredient for models of evolutionary processes. Since variation at the phenotypic level is produced by mutations on the genotypic level, a *genotype-phenotype (GP) map* is needed to model variation quantitatively [2]. Research on GP maps has primarily focused on the simplest case, that of GP maps without non-determinism (ND), where each genotype corresponds to a single, discrete phenotype (Fig. 1). Such GP maps can be characterised by a set of quantitative features [1, 2] (see Fig. 1C, and definitions in table I), which are useful for two reasons: First, some features are sufficiently coarse-grained that they can be estimated from samples even if the full GP map contains too many genotypes to be analysed exhaustively (the number of possible genotypes increases exponentially with sequence length, for example as 4^*L*^ for length-L DNA/RNA sequences) [4, 5]. Secondly, these features can be computed for any given GP map, describing a range of phenotypes at different scales, thus facilitating a more general overview of GP map properties and their evolutionary implications.

**Table 1.**
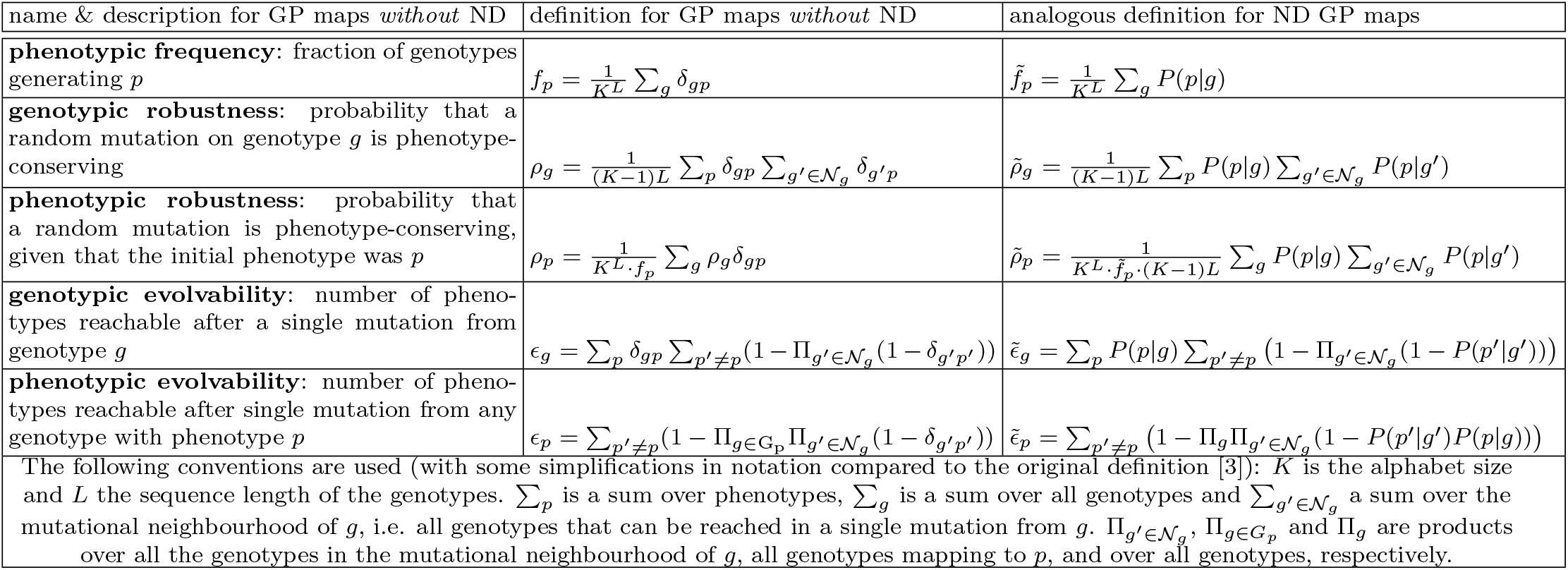
Definitions of commonly used GP map characteristics, both for ‘standard’ GP map without ND and for ND GP maps (see reviews [1, 2] for deterministic case; ref [3] for ND case and notation): The GP map needs to be given as an input (based, for example, on a biophysical model), and is encoded as follows: For GP maps without ND, *δ*_*gp*_ is one if genotype *g* maps to phenotype *p* and zero otherwise. For GP maps with ND, *P*(*p*|*g*) denotes the ensemble probability of phenotype *p* for genotype *g*.

**Figure 1.**
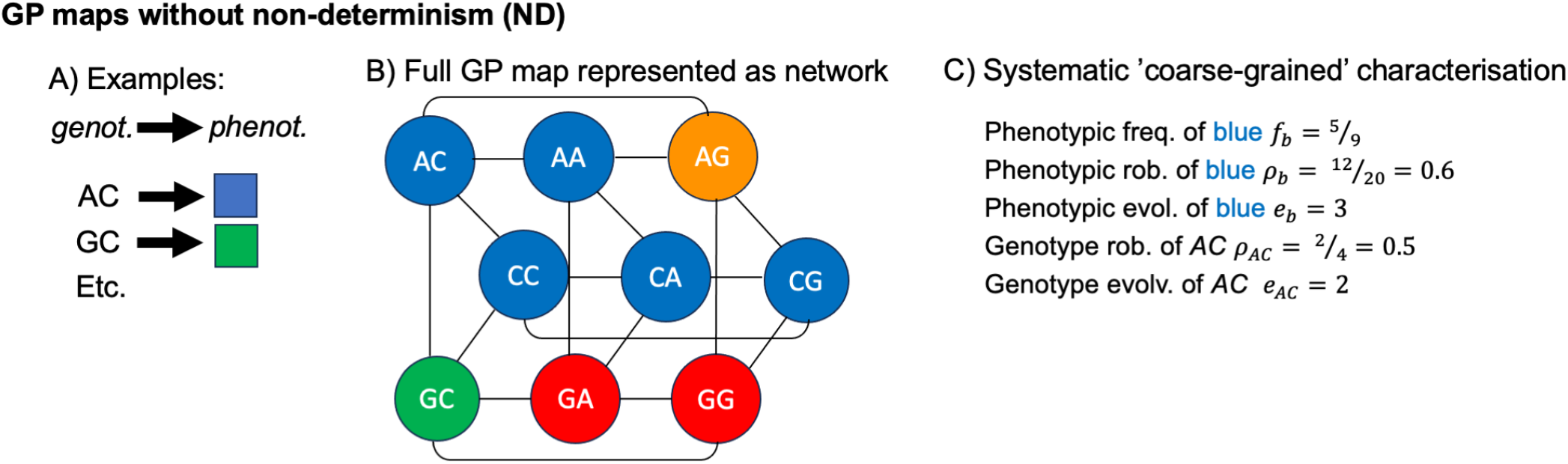
‘Classic’ deterministic GP maps and their characterisation (see reviews [1, 2], and ref [3] for a similar schematic): A) A deterministic GP map is a dataset, where each possible genotype (here sequences of length *L* = 2 and an alphabet size *K* = 3 for simplicity), maps to a single discrete phenotype, here a colour. B) A convenient representation of a GP map is as a network: each genotype is a node, each node has an associated phenotype (here a colour), and edges (grey lines) indicate that two genotypes are only a single point mutation apart. C) Once the GP map is given, it can be characterised with the coarse-grained characteristics defined in *table I*.

Thus, the quantitative features can be computed for different GP maps, describing, for example, protein structures [6], molecular self-assembly phenotypes [7], digital organisms [8, 9] and transcription factor binding [10]. This systematic approach has revealed ‘universal’ characteristics that are found in a range of GP maps (see recent reviews [1, 2]):

1. Some phenotypes are generated by many more genotypes than others (i.e. have different phenotypic frequencies), often by several orders of magnitudes (*phenotypic bias*) [7, 8, 11, 12]. This bias is relevant since high-frequency phenotypes are more likely to appear as variation in evolutionary processes [13, 14].
2. The probability that genotype *g* maps to phenotype *p* is higher if *g* has a mutational neighbour mapping to *p* (*genetic correlations* [15]). These correlations are relevant for neutral evolution, for example [15].
3. A *genotype* that is robust, i.e. has many phenotype-preserving mutations, must be of low evolvability, i.e. only have few distinct phenotypes in its mutational neighbourhood [16]. However, a high-robustness *phenotype* can have high evolvability because phenotypes generated by a large number of genotypes have more mutational neighbours overall [16].

The potential origins of these shared characteristics have been investigated with a toy model, the Fibonacci model [17, 18], which is simple yet sufficient to reproduce these shared characteristics. In this model, each sequence contains a phenotype-dependent set of ‘unconstrained’ positions, which can mutate without phenotypic effect, as well as phenotype-changing ‘constrained’ positions [17]. This concept of phenotype-dependent ‘sequence constraints’ [17] has been used to approximate more complex GP maps [5, 9, 19] and is currently regarded as a central model explaining the origin of the shared characteristics [2]. In addition to this conceptual role, the Fibonacci model is a useful tool for generating GP maps. For example, due to its simplicity, a null model without a positive phenotype-robustness-evolvability relationship can be built [18], which can be used to investigate the relevance of evolvability, for example for the existence of accessible paths [20].

However, the models reviewed so far all ignore one central feature of biological system: in real GP maps, a genotype can produce several phenotypes [23, 24] (Fig. 2). For example, an RNA sequence does not simply fold into a single structure [25], but is better described by an ensemble, where several structures *p* are present in different ratios *P*(*p*|*g*) [26]^1^. Similarly, proteins can have multiple folds [27], self-assembling building blocks can assemble into multiple structures [22], and a single genetic sequence generates a specific distribution of protein sequences due to translational errors [28]. These examples are better captured by ‘non-deterministic’ (ND) GP maps, where each genotype maps to a probability distribution of phenotypes. Such ND [3]/‘plastic’ [29]/‘many-to-many’ [2]/‘probabilistic’ [30] GP maps are fundamentally different from those without ND: Without ND, every genotype corresponds to a single phenotype and thus, every mutation can fall into two classes, either fully phenotype-preserving or generating a new phenotype in a single mutation. With ND, each genotype maps to a phenotype ensemble and mutations can shift the ensemble probabilities by arbitrary amounts.

**Figure 2.**
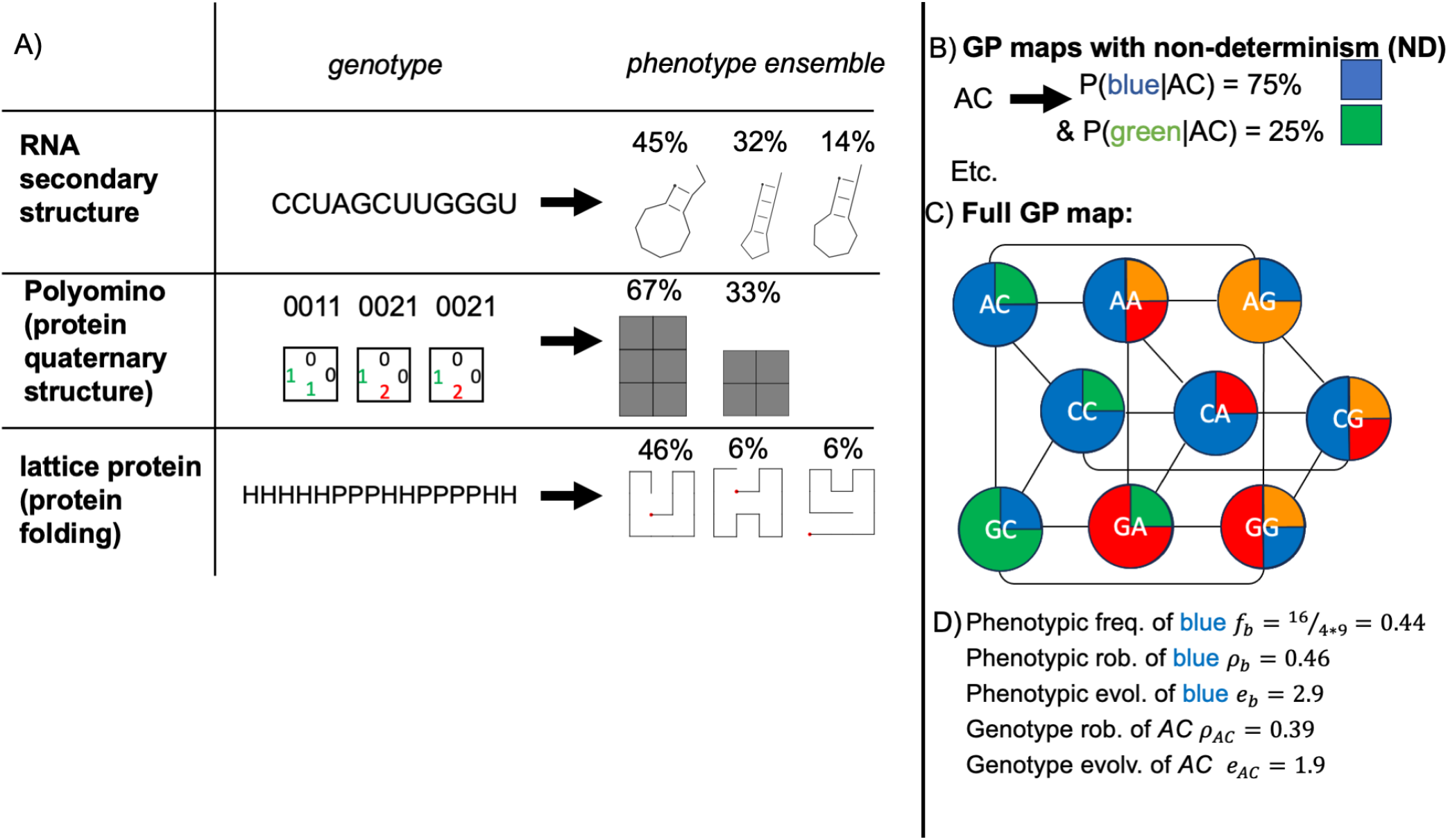
Non-determinism (ND) is present in well-studied biophysical GP maps, and can be included in the GP map framework: **A)** Three well-studied biophysical models, which generate phenotypic information for a given genotype, [1] all contain ND: First is the map from RNA sequences as genotypes to RNA secondary structures as phenotypes [11, 21]. In this model, each sequence can fold into multiple structures p, each with a Boltzmann weight *P*(*p*|*g*). Secondly, in the Polyomino model, a simplified model of protein quaternary structure self-assembly, the genotype is a sequence of integers and defines the binding possibilities of a set of self-assembling 2D tiles (in the pictured example, 1 and 2 bind) [7]. Stochastic self-assemblies with these tiles can give multiple shapes, or phenotypes, each with a probability *P*(*p*|*g*) [22]. The third example is the lattice protein model, a simplified model of protein tertiary structure, where a genotype is an amino acid chain made up of hydrophobic ‘H’ and polar ‘P’ residues [12]. This genotype can fold into multiple structures p, each with a Boltzmann weight *P*(*p*|*g*). Thus, all three models are non-deterministic, and only simplified versions of them fit the deterministic GP map framework (for example, mapping each genotype only to its highest-*P*(*p*|*g*) phenotype). **B)** In their full version, these biophysical models all define ND GP maps, i.e. they map each genotype *g* to several phenotypes *p* and associate a probability *P*(*p*|*g*) to each phenotype. **C)** An ND GP map can be represented as a network (like the deterministic maps in Fig. 1), but now each genotype maps to an ensemble of phenotypes. **D)** Once the ND GP map is given, it can be characterised with the ND analogues of the coarse-grained characteristics, see ref [3] and table I.

ND GP maps are not only more realistic, but also more complex, and thus, it is important to approach them systematically, relying on a consistent set of coarse-grained features. To facilitate this, definitions for GP map features like robustness and evolvability have been proposed for ND GP maps, initially with a threshold-based treatment [22] and later in a treatment that takes full ensemble frequencies into account and proceeds in close analogy with their deterministic counterparts [3, 30] (see table I and Fig. 2). Here, I will focus primarily on the latter definitions, since they consider ensemble frequencies in a more nuanced way, but revisit the former definitions in SI section S5 for completeness. With the second set of definitions, the three ‘universal’ characteristics outlined above were found in the ND version of the RNA GP map [3, 30], and phenotypic bias and genetic correlations also in non-biological input-output maps (spin glasses and quantum circuits) [30]. However, it is not clear, to what extent these features are shared across a wider range of biological GP maps. Moreover, simple models like the Fibonacci model are needed, which put these features into context and can be used to systematically investigate their possible roots, and as a tool to generate simple GP maps for future analyses of GP map features and their implications.

To address this, this paper applies the framework reviewed in table I to two further biophysical ND GP maps (the hydrophobic-polar ‘HP’ lattice model [12], a simple model of protein tertiary structures, and the tile-based ‘Polyomino’ self-assembly model [7] mimicking protein quaternary structure, see Fig. 2). I find that these maps share the ‘universal’ characteristics reviewed above, except the positive correlation between phenotypic robustness and evolvability, which is non-negative, but not always clearly positive. I provide context for these results with a simple *synthetic* GP map. This map shares the ‘universal’ characteristics of the more complex biophysical GP maps, illustrating that few ingredients are needed to define a GP map displaying these shared characteristics.

## II. METHODS

### A. RNA GP map

For each RNA genotype of length *L* = 12 nucleotides, the Boltzmann ensemble of secondary structures was computed using the ViennaRNA package [21] (version 2.7.0): first, a list was generated with all secondary structures whose base pairs are compatible with the genotype *g*. Then, the free energy *G*_*p,g*_ of each structure *p* was calculated with the eval structure function in Vien-naRNA’s Python bindings. Then, the Boltzmann weights follow from the standard relationship:

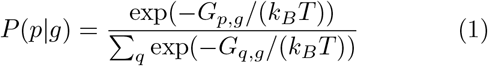

Here, the sum is over all structures *q* that are compatible with *g* (including the unfolded structure with no base pairs).

### B. Lattice protein GP map

In the lattice protein GP map [12, 31], each structure corresponds to a configuration of a polymer chain on a lattice, here a chain of length *L* = 16 on a 4× 4 lattice. To filter for unique configurations, mirror images and rotations were removed (see [32]), but two configurations with reversed chain directions were considered distinct since protein backbones have an inherent directionality. For a given genotype *g*, made up of hydrophobic ‘H’ and polar ‘P’ residues, the free energy *G*_*p,g*_ associated with each configuration *p* was computed with the contact energies in ref [12] (since this HP contact potential gives fewer genotypes with degenerate minimum-energy states than its alternative [32]). Given *G*_*p,g*_, the Boltzmann weights follow from eq. 1. Since the free energy *G*_*p,g*_ is dimensionless in this abstract model, *k*_*B*_*T* is also dimensionless.

### C. Polyomino self-assembly GP map

In the Polyomino model *S*_*t,c*_, a length-4*t* genotype is used to label the 4*t* faces of *t* square tiles, where each tile face can take any integer between 0 and *c* −1 [7]. A face’s integer specifies its binding properties: a face labelled 0 cannot bind, 1 and 2 can bind, 2 with 3 etc. [7].

To go from a genotype - and its encoded set of tiles - to an ensemble of phenotypes, stochastic self-assembly was simulated (using code tested against ref [33]): First, a randomly selected tile from the set initialises the assembly. Then, at each iteration, one tile type that can bind to a free face on the assembly is selected at random and added to the assembly. This process terminates either when there are no options left, or when the assembly exceeds a size threshold ((4*t*)^2^/2 for *t* tiles as in [33]).

Due to the randomness in the process, a single genotype can produce different assemblies (i.e. phenotypes) [33]. To estimate their frequencies, the self-assembly simulation was repeated 500 times per genotype. When processing the outputs, two phenotypes were considered identical if they are rotations or translations of one another (but not mirror images [7]). Further, a single ‘undefined’ placeholder phenotype was used for assemblies exceeding the size threshold, as well as assemblies appearing < 5 times in the assembly process, since their ensemble frequencies cannot be estimated reliably. While repeating the analysis with cut-offs of 3 and 10 gave consistent qualitative conclusions (see SI section S2.5), the limited number of 500 repetitions is a caveat, especially for the characterisation of low-frequency phenotypes.

To reduce computational costs, the assembly graph formalism was used: an assembly graph represents the binding properties of a set of tiles, such that two genotypes with the same assembly graph produce the same ensemble of phenotypes [33]. Thus, assembly was simulated only for one genotype per assembly graph. Genotypes with the same assembly graph were identified in two ways: First, by checking if the assembly-graph-preserving minimal genotype [34] was already attributed to an assembly graph. Secondly, to test whether a given assembly graph matches an already known assembly graph, known assembly graphs were filtered by the number of edges (both in total and per tile) and then NetworkX’s [35] graph isomorphism test was applied to potential matches. While the assembly graph formalism makes a full analysis of the ND GP map computationally feasible, it comes with the caveat that sampling errors in a genotype’s ensemble frequencies are propagated to further genotypes with the same assembly graph.

### D. ‘Undefined’ phenotypes and zero values in the ND GP map analysis

Just as in previous deterministic GP maps [1], an ‘undefined’ phenotype exists in several ND GP map models: an unfolded RNA chain in the RNA model and an ‘undefined’ placeholder in the Polyomino model. Furthermore, the ensemble of the genotype (0, 0, 0,..) in the synthetic model was set to an undefined phenotype, to avoid artefacts (all phenotypes would have a probability 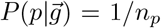 for this genotype). Moreover, in the deterministic version of the synthetic GP map, which maps each genotype 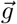 to the lowest-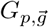 phenotype, the undefined phenotype was used in the case of ties, i.e. when the two lowest-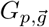 phenotypes differ by less than 10^−4^, in line with conventions for ties in the lattice protein model [15]. These ‘undefined’ structures were not included in the GP map characterisation since they are thought to be artefacts: for RNA, the high prevalence of the unfolded structure is due to the short sequence length of *L* = 12 [1], and in the Polyomino self-assembly, the unfolded structure is a placeholder for both unbounded and rare assemblies and should not be treated as a single phenotype. Thus, the sums ∑_*p*_ in table I were taken over all phenotypes except the ‘undefined’ one, and quantities like phenotypic frequency and robustness were not computed for the ‘undefined’ phenotype.

Further, zeros (for example, 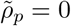 phenotypes) are not shown in log-scale plots.

## III. RESULTS

### A. Shared features of biophysical ND GP maps

Let us start by analysing the ND GP map of three biophysical models (see Fig. 2): RNA secondary structure folding [11, 21], the lattice protein model [12], which describes protein tertiary structure folding in a simplified way, and the Polyomino model of self-assembly [7], which is inspired by protein quaternary structure. All three models have served as key examples of GP maps without ND (see reviews [1, 2]). In addition, they have permitted comparisons with evolved structures in databases, for examples of phenotypic frequencies of RNA secondary structures [36, 37] and the symmetries of protein quaternary structures [38]. While the RNA ND GP map had been analysed before [3, 30], it is included here for completeness. In the Polyomino case, the non-determinism has been pointed out [22, 39], but has only been analysed with a threshold-based treatment [22] and by simplification to a deterministic GP map [39].

#### 1. Phenotypic bias in biophysical ND GP maps

Let us start with the phenotypic frequency 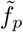, i.e. the mean ensemble frequency of a phenotype *p* across all genotypes and thus its overall abundance (see definition in table I). This frequency of a phenotype *p* is plotted against its rank in each GP map, i.e. against the position of *p* in a list sorted by frequency (Fig. 3, first column). These phenotypic frequencies show phenotypic bias in all three models, in agreement with previous work for RNA [3] and lattice proteins [40]: different phenotypes in a single GP map have different 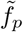. In the RNA and Polyomino models, these frequency differences span several orders of magnitude, and the majority of phenotypes have low 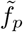 of less than one percent of the maximum phenotypic frequency in the map (for RNA, between 71% and 75%; for Polyomino, between 69% and 84%). In the lattice protein model, the bias is weaker and even the smallest frequency is higher than 0.01 × the largest phe notypic frequency.

**Figure 3.**
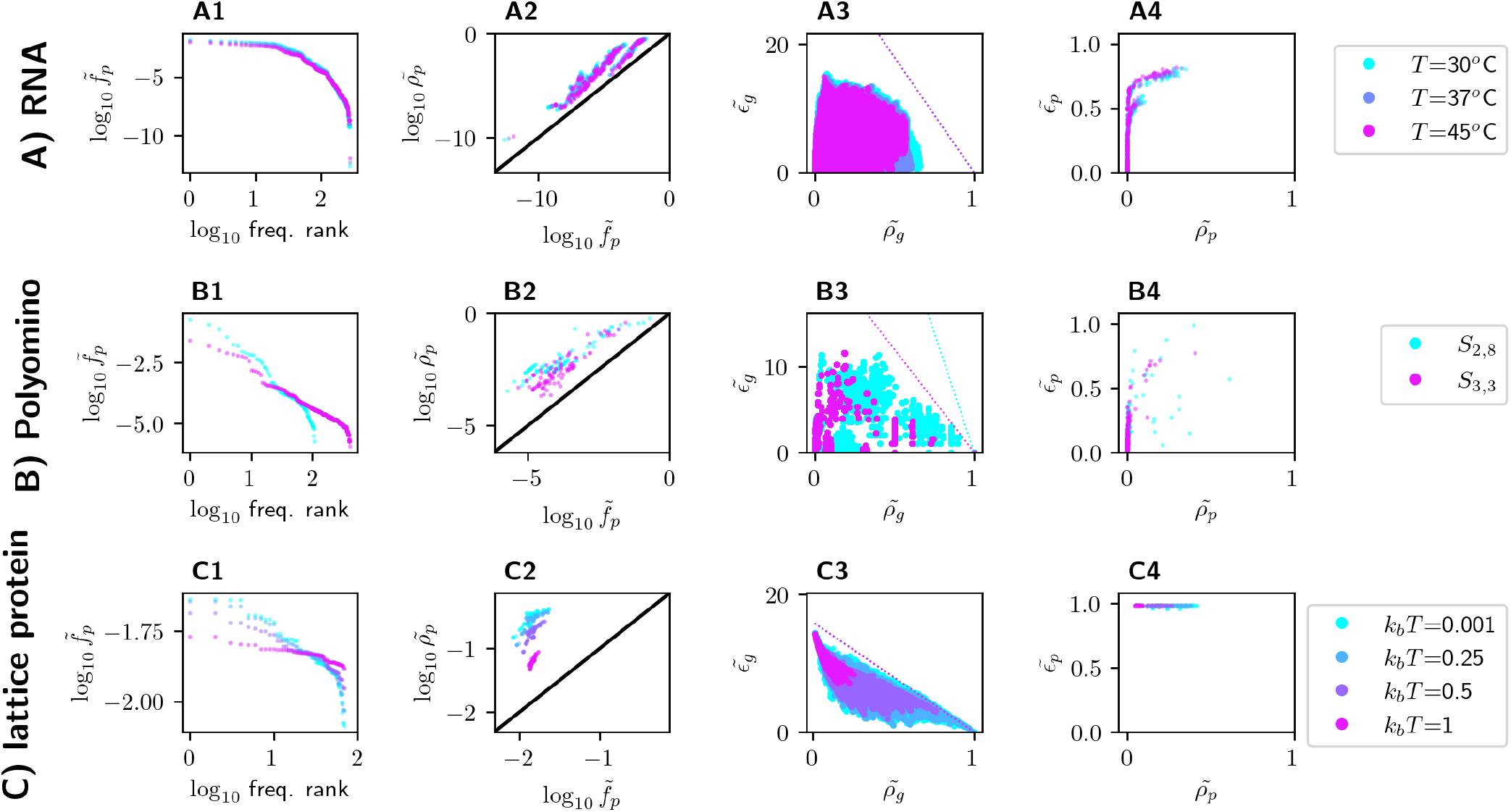
The three biophysical ND GP maps have shared characteristics: Rows **(A) - (C)** show one model each: RNA secondary structure (row **A**, for different folding temperatures; this map had been analysed [3, 30] and is included for completeness), the Polyomino self-assembly model (row **B**, for two parameters, see section II C) and the lattice protein model (row **C**, for different folding temperatures *k*_*B*_*T*). For each ND GP map, the following analyses are shown in columns **(1-4)**:**(1)** The phenotypic frequency 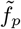 is plotted against the frequency rank, i.e. the position of *p* in a list sorted by frequency, showing phenotypic bias in all cases. **(2)** The phenotypic robustness 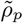 is plotted against the phenotypic frequency 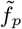 (with *x* = *y* shown as a black line). Since 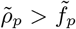 except for zero-robustness phenotypes in the Polyomino *S*_3,3_ case, the maps have genetic correlations. **(3)** The genotypic evolvability 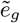 is plotted against genotypic robustness 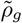. High genotypic evolvability is incompatible with high robustness, consistent with the upper bound *e*_*g*_ < (*K* − 1) *L*(1 − *ρ*_*g*_) (dashed line; the data is below the bound when accounting for numeric errors of up to 10^−4^). **(4)** Phenotypic evolvability 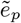 is normalised relative to its maximum *n*_*p*_− 1 and plotted against phenotypic robustness 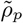 The results are mixed, ranging from RNA with a positive saturating trend to the lattice protein map, where all phenotypes have an evolvability within 5% of the maximum possible value of 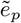.

Looking beyond these commonalities, however, the three biophysical GP maps show differences in their frequency-rank plots, i.e. their cumulative frequency distributions. Thus, the shape of the frequency distributions is model-dependent (see Fig. S1 of the SI). This model-dependence is consistent with GP maps without ND, where different functional forms have been found to approximate frequency distributions in different models [9, 19, 36].

#### 2. Genetic correlations in biophysical ND GP maps

Next, let us focus on genetic correlations, i.e. whether the robustness 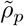 of a phenotype is higher than its phenotypic frequency 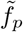. Fig. 3 (second column) shows that this is indeed the case, as previously demonstrated for RNA [3, 30] (more details in Fig. S2 in the SI). The only exception is the the Polyomino *S*_3,3_ model, where 303 out of 398 phenotypes have zero robustness, all low-frequency phenotypes with 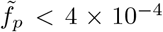. In some of these outliers, the zero robustness may be a numerical consequence of having a fixed minimum *P*(*p*|*g*) ≥ 0.01 (see methods II C), such that neighbours with small, but non-zero *P*(*p*|*g*) are not accounted for: in fact, when lowering the cutoff to 0.006, 68 of these phenotypes show genetic correlations. Because of this and since all high-frequency phenotypes have 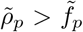, I consider this map to have genetic correlations.

The presence of genetic correlations is further supported by the correlations between ensemble frequencies: in GP maps with genetic correlations, a genotype is more likely to have *p* as a high-probability structure in its ensemble if a neighbouring genotype has *p* as a high-probability structure (see SI section S1.1). Such a correlation is indeed found in the three maps (Fig. S3 in the SI, again with exceptions in the Polyomino case).

#### 3. Genotypic robustness and evolvability in biophysical ND GP maps

Let us now turn to the relationship between the robustness 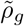 and evolvability 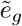 of individual *genotypes*. In the deterministic case, genotypes have a limited number of mutational neighbours and thus cannot combine high robustness with high evolvability, giving (1 − *ρ*_*g*_)(*K* − 1)*L* ≤ *e*_*g*_ [16]. This trade-off generalises to the ND case (SI section S1.2) and is consistent with the data for the three biophysical examples (Fig. 3, third column). Note that the RNA maps contain more low-robustness-low-evolvability genotypes than in a previous analysis [3], likely because I excluded the ‘undefined’ phenotype from the calculations. However, for the qualitative conclusions this does not matter and there is a genotypic robustness-evolvability trade-off in all three maps.

#### 4. Phenotypic robustness and evolvability in biophysical ND GP maps

Despite the trade-off between genotypic evolvability and robustness, the *phenotypic* analogues can be positively correlated. This is simplest to understand for deterministic GP maps: more robust phenotypes have higher phenotypic frequencies, and thus more mutational neighbours overall, giving them the potential for higher evolvability [16]. This trend can continue until the maximum possible evolvability value, which is given by the total number of phenotypes *n*_*p*_ (minus the source phenotype).

When plotting phenotypic evolvability against robustness in the biophysical ND GP maps, the results are mixed (Fig. 3, fourth column): In the RNA map, there is a clear positive trend (in agreement with ref [3]), but in the Polyomino map, this trend is weaker, particularly in the two-tile system S_2,8_ (compared to the three-tile case S_3,3_). Thus, the weak trend may be an artefact of the relatively small system size, chosen for reasons of computational feasibility. In the lattice protein model, all phenotypes have evolvabilities within 5% of the maximum, and thus there is no clear trend. This is reminiscent of 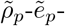 relationships with a saturating trend in deterministic GP maps (of transcription factor binding [41], also a randomised model without genetic correlations [42]), except that the lattice protein ND GP map does not have a single normalised evolvability value below 95%. I will hypothesise possible reasons for this saturation after comparing it to the synthetic ND GP map.

### B. A simple synthetic GP map shares the characteristics of more complex biophysical ND GP maps

The previous section has shown that three biophysical ND GP maps share key features of GP maps without ND: phenotypic bias, genetic correlations, and a robustness/evolvability relationship that is negative on the genotypic and non-negative on the phenotypic level. This prompts the following question: Which ingredients are needed in a model that replicates these features? To answer this, let us consider a ND GP map without any underlying biophysical model and with only three free parameters (Fig. 4).

**Figure 4.**
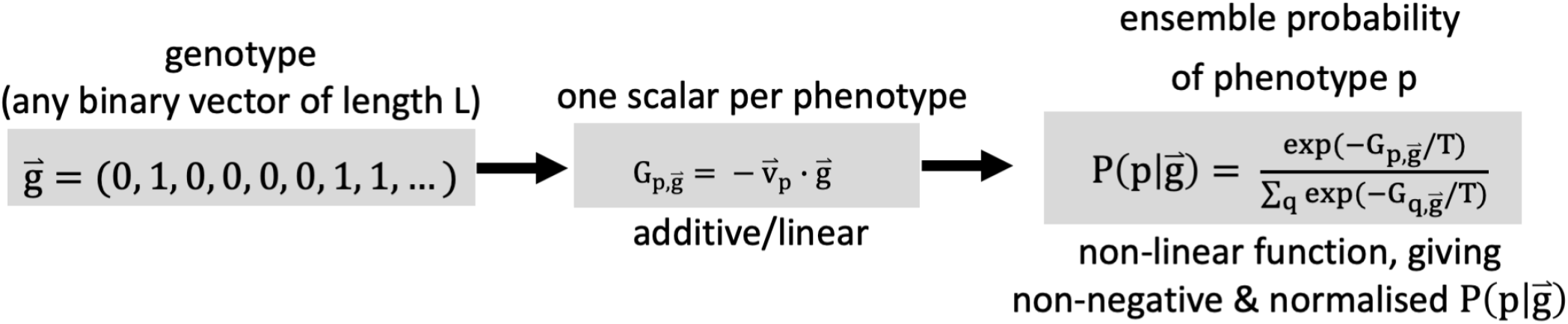
Synthetic ND GP map: each genotype, an arbitrary binary vector of length *L*, is mapped to a distribution of phenotypes. Before constructing the GP map, the following parameters need to be set: the number of phenotypes *n*_*p*_, the stochasticity *T*, the sequence length *L*, and one *L*-dimensional vector 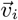 for each phenotype, which is drawn from a Gaussian distribution (with s.d. and mean = 1). The free parameters (*n*_*p*_, *T*, *L*) and the sampled parameters 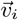 are sufficient to construct the entire ND GP map, by calculating the probability 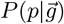 for any of the *n*_*p*_ phenotypes *p* and any of the 2^*L*^ genotypes 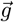.

#### 1. Definition of a simple synthetic ND GP map

To produce a ND GP map, the synthetic model needs to map a genotype 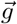, i.e. a sequence of characters from a fixed alphabet, to an ensemble of phenotypes, given by a valid probability distribution (i.e. non-negative ensemble frequencies summing to one). Further, since epistasis is important in evolutionary landscapes [43], the function should contain non-linearities. These objectives are met, for example, by the following Boltzmann-ensemble-inspired function over *n*_*p*_ phenotypes *q*:

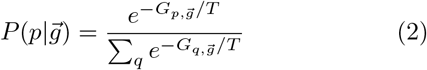

Here, *T* is analogous to the temperature in a Boltzmann ensemble. Thus, varying T from *T* → 0 to *T* → ∞ takes the map from the deterministic limit, which is dominated by a single phenotype per genotype, to the extremely non-deterministic limit, in which all phenotypes have probabilities *P*(*p*|*g*) = 1/*n*_*p*_ for any genotype *g* and the ND GP map is so simple that it can be characterised analytically (SI section S3.2). In fact, in this limit the map becomes an extreme case of the correlation-free null model where *P*(*p*|*g*) = *f*_*p*_ [30].

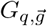 is analogous to the free energy and needs to be a scalar depending on both genotype and phenotype. For simplicity, the model will take binary genotypes from an alphabet of ‘1’ and ‘0’, which can be represented as an *L*-dimensional binary vector 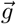. Then, a scalar quantity can be obtained from 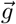 using a linear, additive function, which can be written as a dot product:

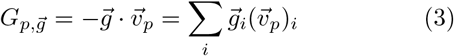

Here, the phenotype-dependent, *L*-dimensional vector 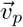 contains the parameters of the linear function. It only needs to be initialised once for each phenotype *p* and is then used for all 2^*L*^ genotypes in the map. To avoid setting additional parameters, each element of 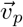 is drawn from a normal distribution (similar to existing examples of GP maps/fitness landscapes with randomly drawn parameters [44, 45]). The mean and standard deviation of the normal distribution are simply taken to be one since any scaling factors would cancel in the normalisation or by adjusting *T* (similar to [45]).

These steps define a synthetic ND GP map with only three parameters to choose (sequence length *L*, number of phenotypes *n*_*p*_ and stochasticity *T*). The ND GP map closely resembles a recent two-phenotype model for bottlenecks in mutational paths [45], which also combines a Boltzmann-like nonlinearity with a linear function and randomly sampled coefficients. However, this paper assumes more than two phenotypes, and focuses on a different question: whether this model reproduces the shared ND GP map properties. This analysis will consider several choices of *n*_*p*_ (first row of Fig. 5) and T (second row of Fig. 5), for a fixed sequence length *L* = 15.

**Figure 5.**
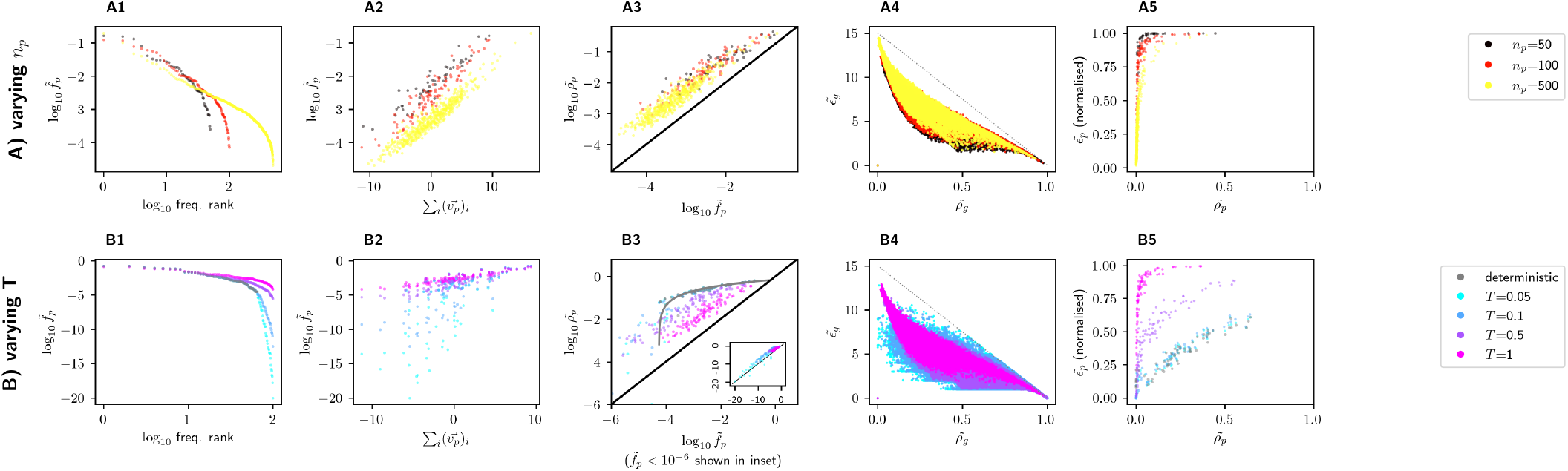
The synthetic ND GP map replicates the shared features of the biophysical ND GP maps: Row **A** shows maps with different numbers of phenotypes *n*_*p*_ (to maintain a comparable level of non-determinism, *T* is set to the genotypic average of the 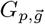-difference between the lowest two phenotypes), and row **B** focuses on maps with different stochasticity *T* for fixed *n*_*p*_ = 100 (including the deterministic limit *T* →0, where each genotype maps to the lowest-G structure). In both rows, the following analyses are shown in columns **(1) - (5)**: **(1)** The phenotypic frequency 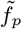 is plotted against the frequency rank, i.e. the position of *p* in a list sorted by frequency, showing phenotypic bias. **(2)** The phenotypic frequency 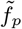 is plotted against the element sum of the corresponding parameter vector 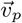. The positive trend is consistent with the hypothesis that small differences in the parameter vectors of different phenotypes *p* are responsible for their different phenotypic frequencies 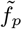. **(3)** The phenotypic robustness 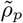 is plotted against the phenotypic frequency 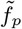 (with *x* = *y* indicated by a black line).Since 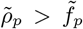 for the majority of phenotypes, there are genetic correlations. In B3, only phenotypes with 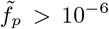are shown; the same plot with the full range of 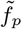 is included as an inset. **(4)** Genotypic evolvability 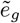 is plotted against genotypic robustness 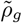. There are no high-evolvability-high-robustness genotypes, consistent with the upper bound *e*_*g*_ < (*K−*1)*L*(1*−ρ*_*g*_) (dashed line; the data is below the bound when accounting for numeric errors of up to 10^−4^). **(5)** Phenotypic evolvability 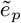 is normalised relative to its maximum *n*_*p*_ −1^a^ and plotted against phenotypic robustness 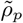, showing a positive trend with saturation in some maps. ^a^ Here, the *n*_*p*_ in the map setup is used for the normalisation, even though some phenotypes may not appear in the map, especially in the deterministic limit.

#### 2. Phenotypic bias in the synthetic ND GP map

Let us start with phenotypic frequencies 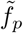 in the synthetic map (first column of Fig. 5): phenotypic frequencies in the map can range over several orders of magnitude, i.e. the synthetic model has phenotypic bias. Since phenotypes only differ in their parameters 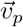, this bias must emerge from random differences in 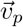. Concretely, let us consider the following hypothesis: a vector with larger elements tends to give more favourable values of 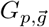, which are then amplified by the non-linear function in eq. 2 and summed over all genotypes, leading to the observed frequency differences of several orders of magnitude. To test this hypothesis, the element sum of 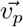 is plotted against 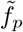 for each phenotype *p* in the second column of Fig. 5, giving a positive trend as hypothesised. This argument is reminiscent of research on the deterministic GP maps of RNA and lattice proteins [46, 47], in that it links the free energy differences for individual genotypes to differences in phenotypic frequencies in the entire GP map.Comparing maps constructed with different parameters, we find phenotypic bias to weaken with increasing stochasticity *T*. A higher stochasticity *T* reduces the impact of phenotypic differences in 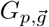 on ensemble probabilities for each individual genotype (see eq. 2). Thus, phenotypic differences will also be less pronounced when summed over all genotypes to obtain phenotypic frequencies 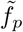. In the extreme case *T* → ∞, phenotypic differences vanish entirely, giving 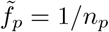 for all phenotypes (SI section S3.2) and thus no phenotypic bias.

Further, the number of phenotypes is lower in the fully deterministic limit, where each genotype maps to its lowest-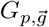 phenotype (65 instead of 100), as is the case in the biophysical RNA map [3]: in this limit, phenotypes only appear if they are the lowest-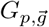 phenotype for one or more genotypes.

#### 3. Genetic correlations in the synthetic ND GP map

Let us now turn to genetic correlation, i.e. whether the robustness 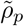 of a phenotype is typically higher than its frequency 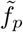. This is indeed the case (Fig. 5, third column), but with exceptions among low-frequency phenotypes with frequencies lower than 3.1 ×10^−5^, which is on the same order of magnitude as the the minimum possible frequency in the deterministic GP map 1/*K*^*L*^ (see inset of Fig. 5 B3). In the synthetic model, the root of these genetic correlations is simple to understand: a single mutation only changes a single site of the genotype and thus a single summand in the dot product of eq. 3. Thus, if phenotype *p* has low 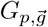 and high ensemble probability for genotype 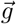, *p* is also likely to have low 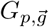 and high ensemble probability for genotypes 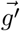 that are one point mutation away from 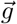.

When comparing the ND GP map with different stochasticities *T* (Fig. 5 B3), we find that a higher stochasticity tends to weaken genetic correlations. This is best understood in the extreme limit *T* → ∞, where *P*(*p*|*g*) = 1/*n*_*p*_ for all phenotypes and genotypes, and thus 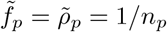,for all phenotypes (SI section S3.2). Thus, in the *T* → ∞ limit, there is no local, genotype-dependent structure and no genetic correlations.

Focussing on the shape of the frequency-robustness relationship in more detail, the well-studied [48] log-linear frequency-robustness scaling provides a good fit in the deterministic limit (grey line in Fig. 5 B3). In the ND case, the data deviates from the deterministic scaling, consistent with the biophysical maps and previous analyses of the RNA map as well as quantum circuits and spin glasses [3, 30]. We find that 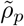 in ND GP maps depends not just on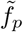, consistent with a recent scaling expressing 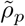 in terms of two quantities: 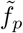 and 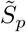, which can depend on *T* and describes the number of genotypes a phenotype is concentrated on [30] (see SI section S3.1).

#### 4. Genotypic robustness and evolvability in the synthetic ND GP map

Turning to the relationship between *genotypic* robustness and evolvability, Fig. 5 (fourth column) shows that there are no high-robustness-high-evolvability genotypes, in agreement with the upper bound. While some maps have both high-robustness-low-evolvability and low-robustness-high-evolvability genotypes, high-*T* maps are limited to low-robustness genotypes. Again, this can be understood by considering the extreme limit *T* → ∞ with 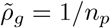 and 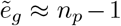 for all genotypes (SI section S3.2).

#### 5. Phenotypic robustness and evolvability in the synthetic ND GP map

Finally, the fifth column of Fig. 5 shows 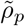 and 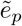 on the *phenotypic* level. Higher-robustness phenotypes tend to be more evolvable, with a potential saturation at the maximum possible evolvability, *n*_*p*_ − 1. This saturation is especially pronounced in two parameter regimes, a low number of phenotypes *n*_*p*_ or high stochasticity *T*: For GP maps with a low number of phenotypes *n*_*p*_ − 1, the maximum possible evolvability value 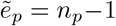 is low and therefore reached faster. In GP maps with high stochasticity *T*, the phenotypic diversity in any mutational neighbourhood is high (demonstrated by the high *genotypic* evolvabilities), leading to high phenotypic evolvability values and thus to saturation. In the extreme limit *T* → ∞, all phenotypes have 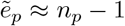 (SI section S3.2).

#### 6. Alternative definitions of the synthetic ND GP map

The synthetic model displays the shared ND GP map characteristics for a range of *n*_*p*_ and *T*, so the next question is, whether these results depend strongly on the other choices made when designing the model: the Boltzmann-like non-linear function in the GP map, the normal distribution used to initialise the parameters, and the binary genotype alphabet. Thus, I built synthetic ND GP maps with three alternative non-linear functions (with a shifted linear, a shifted inverse-squared and a shifted Gaussian in the numerator), three alternative distributions for parameter initialisation (uniform/lognormal/exponential distributions), and a different alphabet size of *K* = 4. These alternative ND GP maps continue to reproduce the shared features (see SI section S4). However, the model built with a shifted linear function is an outlier with properties similar to the high-*T* limit: less than an order of magnitude difference between the highest and lowest phenotypic frequencies, weak genetic correlations, low genotypic robustness throughout the map, and phenotypic evolvabilities saturated at the maximum *n*_*p*_−1. This parallel is not surprising since both the linear model and the high-*T* limit do not strongly suppress the ensemble frequencies of a genotype’s less favoured, high-G phenotypes. In other words, both cases are closer to the trivial null model with *P*(*p*|*g*) = 1/*n*_*p*_ for each genotype *g* and phenotype *p*. Thus, some mechanism biasing phenotypic ensembles, such as non-linearities, may be a key part of realistic GP maps, enabling strong bias and genetic correlations. Despite this outlier, there are various versions of the synthetic model reproducing the shared features of the biophysical maps, suggesting that these features may emerge easily from simple models.

## IV. DISCUSSION AND CONCLUSIONS

While many realistic genotype-phenotype (GP) maps are non-deterministic (ND), our understanding of such maps is much less developed than that of simpler GP maps without ND, where every genotype maps to a single categorical phenotype. Here, I fill this gap both by analysing biophysical ND GP maps systematically, following a recent framework [3], and by providing context with a synthetic ND GP map.

The analysis of three biophysical GP maps - the RNA secondary structure model (previously analysed in [3, 30]), the lattice protein model for protein tertiary structure and the Polyomino model for protein quaternary structure - indicates that key ‘universal’ properties of GP maps without ND continue to hold: phenotypic bias, genetic correlations, and a negative relationship between genotypic evolvability and robustness that becomes non-negative on the phenotypic level.

Next, I put these shared properties into context with a synthetic ND GP map. This map, as well as variations with different functional forms and different initialisations, reproduce the shared features of the biophysical models, indicating that they emerge from simple, non-biological models without many ingredients or finetuning, and may thus be found more widely.

The synthetic ND GP map combines an additive genotype-dependence (eq. 3) with a non-linear scalar function (eq. 2). Replacing the non-linear function with a normalised and scaled linear function gave only weak phenotypic bias and genetic correlations, suggesting that mechanisms biasing phenotypic ensembles, such as non-linearities, may be important in realistic models. The combination of additive scalars with a non-linear scalar function is reminiscent of fits to empirical protein sequence-function/fitness data [49–51], and highly similar to a recent two-phenotype model, which captured bottlenecks in mutational paths and was also initialised with randomly drawn parameters [45]. Note, however, that the multi-phenotype case differs from the two-phenotype case [52], which can be still written in terms of a single genotype-dependent scalar 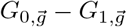 [45].

Interestingly, the high-stochasticity limit of the synthetic ND GP map shares features with the lattice protein model, e.g. the saturated phenotypic evolvabilities and weak phenotypic bias. These parallels may stem from the high stochasticity in the lattice protein model: ≈ 62% of lattice protein genotypes have more than one minimum-energy structure and thus even the highest ensemble frequency cannot reach 50%.

In its deterministic limit, the synthetic model replicates the well-studied properties of deterministic GP maps. These shared properties had previously been explained with constraint-based models, but the synthetic model differs from idealised constraint-based models like the Fibonacci model: the availability of phenotype-conserving, neutral mutations at a given position is highly sequence-dependent (more details in SI, section S3.3). This sequence-dependence is interesting, since there is some evidence that it also exists in (models of) biological systems: an empirical GP map for transcription factor binding (here the sequence-dependence is seen in the marked variation in genotypic robustness within a phenotype) [41], a Potts-model-fit to the *β*-lactamase sequence family [53] and empirical fitness effects in orthologous sequences [54]. Future work should investigate these sequence-dependent effects in more detail, and whether they are sufficiently strong for these examples to fall outside the scope of ‘constraint-based’ models.

Since the synthetic model replicates key features of biophysical models, it can serve as a tractable model for future work on ND GP maps and their implications for evolutionary processes: The model allows the construction of ND GP maps of arbitrary size, by flexibly choosing the sequence length *L* and number of phenotypes *n*_*p*_.

This flexibility is not available in biophysical models: for example, in the RNA model, a sequence length of L = 12 is necessary to have 273 non-trivial phenotypes - and thus 4^12^ ≈ 1.7 × 10^7^ genotypes. In addition to this flexibility in setting the number of genotypes and phenotypes, the synthetic model has a tuneable stochasticity T and can be extended to larger alphabet sizes (SI section S4.3), allowing us to generate a family of ND GP maps whose features can be contrasted. Thus, it can be used in a similar way as the Fibonacci model, where the contrast between a low- and a high-evolvability version [18] has been used to test the role of evolvability in permitting accessible paths [20].

Just as statistical ‘House-of-Cards’/’NK’/’Rough-Mount-Fuji’ models are used to address questions about fitness landscapes [55], the synthetic model may thus help investigate ND GP maps systematically, in particular:

- **Developing sampling methods:** Further developments of sampling methods (see [3]) could allow estimates of ND GP map features (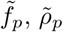 ideally even 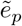), from small and local samples. Then, these features could be estimated for more complex computational models, e.g. gene regulatory networks [56], and from high-throughput experimental data.
- **Evolutionary simulations:** Quantities like phenotypic evolvability were motivated by deterministic GP maps, where population can drift through the set of phenotypically identical genotypes [16]. However, in ND GP maps, genotypes dominated by the same phenotype *p* typically have different ensemble probabilities *P*(*p*|*g*) (compare Figs. 1 & 2) and thus the notion of phenotypically identical genotypes no longer applies [29]. Thus, instead of evolvability and other analogues from deterministic maps, new coarse-grained quantities should be defined to quantify aspects of ND GP maps relevant under different evolutionary scenarios (e.g. changing environments [57], selection for multiple phenotypes [58]).

Since ND GP maps combine discrete and continuos phenotypic information (discrete phenotypes *p* and continuous probabilities *P*(*p*|*g*)), the evolutionary simulations could be generalised to a related class of GP maps, where *P*(*p*|*g*) are no longer probabilities. This would include a wider range of systems: for example, empirical transcription factor binding and RNA-binding protein GP maps [59, 60], where each sequence maps to a set of discrete transcription factors/RNA binding proteins, as well as a continuous enrichment score for each transcription factor/RNA binding protein.

## V. ACKNOWLEDGEMENTS

I acknowledge support of the Spanish Ministry of Science and Innovation through the Centro de Excelencia Severo Ochoa (CEX2020-001049-S, MCIN/AEI/10.13039/501100011033), the EMBL partnership and the Generalitat de Catalunya through the CERCA programme. This research is part of Grant JDC2022-049526-I funded by MCIN/AEI/10.13039/501100011033 and by European Union NextGenerationEU/PRTR. I thank M. Giraud for helpful comments.

## VI. DATA AVAILABILITY

The code behind this analysis can be found at https://github.com/noramartin/simple_models.

## Footnotes

1 Here, this ensemble will be a Boltzmann ensemble for computational feasibility, ignoring RNA modifications, kinetic effects, alternative splicing etc., but for the choice of a GP map framework, what matters is that realistic GP maps go beyond the sequence-to-single-categorical-structure paradigm.

## Notes

### Competing Interest Statement

The authors have declared no competing interest.

## References

[1] S. E. Ahnert, Structural properties of genotype–phenotype maps, J. R. Soc. Interface 14, 20170275 (2017).

[2] S. Manrubia, J. A. Cuesta, J. Aguirre, S. E. Ahnert, L. Altenberg, A. V. Cano, et al., From genotypes to organisms: State-of-the-art and perspectives of a cornerstone in evolutionary dynamics, Physics of Life Reviews 38, 55 (2021).

[3] P. García-Galindo, S. E. Ahnert, and N. S. Martin, The non-deterministic genotype–phenotype map of RNA secondary structure, J. R. Soc. Interface 20, 20230132 (2023).

[4] T. Jörg, O. C. Martin, and A. Wagner, Neutral network sizes of biological RNA molecules can be computed and are not atypically small, BMC bioinformatics 9, 464 (2008).

[5] M. Weiß and S. E. Ahnert, Using small samples to estimate neutral component size and robustness in the genotype–phenotype map of RNA secondary structure, J. R. Soc. Interface 17, 20190784 (2020).

[6] E. Ferrada and A. Wagner, A comparison of genotypephenotype maps for RNA and proteins, Biophys. J. 102, 1916 (2012).

[7] S. F. Greenbury, I. G. Johnston, A. A. Louis, and S. E. Ahnert, A tractable genotype–phenotype map modelling the self-assembly of protein quaternary structure, J. R. Soc. Interface 11, 20140249 (2014).

[8] M. A. Fortuna, L. Zaman, C. Ofria, and A. Wagner, The genotype-phenotype map of an evolving digital organism, PLOS Comput. Biol. 13, e1005414 (2017).

[9] N. S. Martin, C. Q. Camargo, and A. A. Louis, Bias in the arrival of variation can dominate over natural selection in Richard Dawkins’s biomorphs, PLOS Comput. Biol. 20, e1011893 (2024).

[10] J. Aguilar-Rodríguez, L. Peel, M. Stella, A. Wagner, and J. L. Payne, The architecture of an empirical genotype-phenotype map, Evolution 72, 1242 (2018).

[11] P. Schuster, W. Fontana, P. F. Stadler, and I. L. Hofacker, From sequences to shapes and back: a case study in RNA secondary structures, Proc. R. Soc. Lond. B 255, 279 (1994).

[12] H. Li, R. Helling, C. Tang, and N. Wingreen, Emergence of preferred structures in a simple model of protein folding, Science 273, 666 (1996).

[13] M. C. Cowperthwaite, E. P. Economo, W. R. Harcombe, E. L. Miller, and L. A. Meyers, The ascent of the abundant: how mutational networks constrain evolution, PLOS Comput. Biol. 4, e1000110 (2008).

[14] S. Schaper and A. A. Louis, The arrival of the frequent: how bias in genotype-phenotype maps can steer populations to local optima, PloS one 9, e86635 (2014).

[15] S. F. Greenbury, S. Schaper, S. E. Ahnert, and A. A. Louis, Genetic correlations greatly increase mutational robustness and can both reduce and enhance evolvability, PLOS Comput. Biol. 12, e1004773 (2016).

[16] A. Wagner, Robustness and evolvability: a paradox resolved, Proc. R. Soc. Lond. B 275, 91 (2008).

[17] S. Greenbury and S. E. Ahnert, The organization of biological sequences into constrained and unconstrained parts determines fundamental properties of genotype– phenotype maps, J. R. Soc. Interface 12, 20150724 (2015).

[18] M. Weiß and S. E. Ahnert, Phenotypes can be robust and evolvable if mutations have non-local effects on sequence constraints, J. R. Soc. Interface 15, 20170618 (2018).

[19] J. A. García-Martín, P. Catalán, S. Manrubia, and J. A. Cuesta, Statistical theory of phenotype abundance distributions: A test through exact enumeration of genotype spaces, EPL 123, 28001 (2018).

[20] M. Srivastava, A. A. Louis, and N. S. Martin, Predicting the topography of fitness landscapes from the structure of genotype-phenotype maps, bioRxiv, 2025.02.14.638275 (2025).

[21] R. Lorenz, S. H. Bernhart, C. Höner zu Siederdissen, H. Tafer, C. Flamm, P. F. Stadler, and I. L. Hofacker, Viennarna package 2.0, Algorithms for molecular biology 6, 1 (2011).

[22] V. Jouffrey, A. Leonard, and S. Ahnert, Gene duplication and subsequent diversification strongly affect phenotypic evolvability and robustness, R. Soc. Open Sci. 8, 201636 (2021).

[23] D. S. Tawfik, Messy biology and the origins of evolutionary innovations, Nature chemical biology 6, 692 (2010).

[24] D. P. Bendixsen, J. Collet, B. Østman, and E. J. Hayden, Genotype network intersections promote evolutionary innovation, PLoS Biology 17, e3000300 (2019).

[25] R. Bose, I. Saleem, and A. M. Mustoe, Causes, functions, and therapeutic possibilities of RNA secondary structure ensembles and alternative states, Cell Chemical Biology 31, 17 (2024).

[26] G. Steger and R. Giegerich, 14. RNA structure prediction, in RNA Structure and Folding, edited by D. Klostermeier and C. Hammann (De Gruyter, 2013) pp. 335–362.

[27] L. L. Porter and L. L. Looger, Extant fold-switching proteins are widespread, PNAS 115, 5968 (2018).

[28] C. Landerer, J. Poehls, and A. Toth-Petroczy, Fitness Effects of Phenotypic Mutations at Proteome-Scale Reveal Optimality of Translation Machinery, Mol. Biol. Evol. 41, msae048 (2024).

[29] L. W. Ancel and W. Fontana, Plasticity, evolvability, and modularity in RNA, Journal of Experimental Zoology 288, 242 (2000).

[30] A. Sappington and V. Mohanty, Probabilistic genotype-phenotype maps reveal mutational robustness of RNA folding, spin glasses, and quantum circuits, Phys. Rev. Research 7, 013118 (2025).

[31] K. F. Lau and K. A. Dill, A lattice statistical mechanics model of the conformational and sequence spaces of proteins, Macromolecules 22, 3986 (1989).

[32] N. E. Buchler and R. A. Goldstein, Effect of alphabet size and foldability requirements on protein structure designability, Proteins: Structure, Function, and Bioinformatics 34, 113 (1999).

[33] S. Tesoro, S. Ahnert, and A. Leonard, Determinism and boundedness of self-assembling structures, Phys. Rev. E 98, 022113 (2018).

[34] A. Leonard, Modelling the evolution of biological complexity with a two-dimensional lattice self-assembly process, Ph.D. thesis, University of Cambridge (2020).

[35] A. Hagberg, P. J. Swart, and D. A. Schult, Exploring network structure, dynamics, and function using NetworkX, Tech. Rep. (Los Alamos National Laboratory (LANL), Los Alamos, NM (United States), 2008).

[36] K. Dingle, S. Schaper, and A. A. Louis, The structure of the genotype–phenotype map strongly constrains the evolution of non-coding RNA, Interface focus 5, 20150053 (2015).

[37] K. Dingle, F. Ghaddar, P. Šulc, and A. A. Louis, Phenotype bias determines how natural RNA structures occupy the morphospace of all possible shapes, Mol. Biol. Evol. 39, msab280 (2022).

[38] I. G. Johnston, K. Dingle, S. F. Greenbury, C. Q. Camargo, J. P. Doye, S. E. Ahnert, and A. A. Louis, Symmetry and simplicity spontaneously emerge from the algorithmic nature of evolution, PNAS 119, e2113883119 (2022).

[39] S. Tesoro and S. Ahnert, Non-deterministic genotype-phenotype maps of biological self-assembly, Europhysics Letters 123, 38002 (2018).

[40] N. S. Martin and S. E. Ahnert, The Boltzmann distributions of molecular structures predict likely changes through random mutations, Biophys. J. 122, 4467 (2023).

[41] J. L. Payne and A. Wagner, The robustness and evolvability of transcription factor binding sites, Science 343, 875 (2014).

[42] X. Wei and J. Zhang, Why phenotype robustness promotes phenotype evolvability, Genome Biol. Evol 9, 3509 (2017).

[43] J. Domingo, P. Baeza-Centurion, and B. Lehner, The causes and consequences of genetic interactions (epistasis), Annu. Rev. Genom. Hum. Genet. 20, 433 (2019).

[44] S. G. Das, M. Mungan, and J. Krug, Epistasis-mediated compensatory evolution in a fitness landscape with adaptational tradeoffs, PNAS 122, e2422520122 (2025).

[45] A. Ottavia Schulte, S. Alqatari, S. Rossi, and F. Zamponi, Functional bottlenecks can emerge from non-epistatic underlying traits, bioarXiv, 2025.05.20.655048 (2025).

[46] N. S. Martin and S. E. Ahnert, Fast free-energy-based neutral set size estimates for the RNA genotype– phenotype map, J. R. Soc. Interface 19, 20220072 (2022).

[47] J. L. England and E. I. Shakhnovich, Structural determinant of protein designability, PRL 90, 218101 (2003).

[48] V. Mohanty, Robustness of evolutionary and glassy systems, Ph.D. thesis, University of Oxford (2021).

[49] J. Otwinowski, D. M. McCandlish, and J. B. Plotkin, Inferring the shape of global epistasis, PNAS 115, E7550 (2018).

[50] K. E. Johansson, K. Lindorff-Larsen, and J. R. Winther, Global analysis of multi-mutants to improve protein function, J. Mol. Biol 435, 168034 (2023).

[51] P. D. Tonner, A. Pressman, and D. Ross, Interpretable modeling of genotype–phenotype landscapes with state-of-the-art predictive power, PNAS 119, e2114021119 (2022).

[52] A. J. Morrison, D. R. Wonderlick, and M. J. Harms, Ensemble epistasis: thermodynamic origins of nonadditivity between mutations, Genetics 219, iyab105 (2021).

[53] L. Di Bari, M. Bisardi, S. Cotogno, M. Weigt, and F. Zamponi, Emergent time scales of epistasis in protein evolution, PNAS 121, e2406807121 (2024).

[54] V. O. Pokusaeva, D. R. Usmanova, E. V. Putintseva, L. Espinar, K. S. Sarkisyan, A. S. Mishin, et al., An experimental assay of the interactions of amino acids from orthologous sequences shaping a complex fitness landscape, PLoS Genet. 15, e1008079 (2019).

[55] I. Fragata, A. Blanckaert, M. A. D. Louro, D. A. Liberles, and C. Bank, Evolution in the light of fitness landscape theory, Trends in ecology & evolution 34, 69 (2019).

[56] C. Espinosa-Soto, O. C. Martin, and A. Wagner, Phenotypic plasticity can facilitate adaptive evolution in gene regulatory circuits, BMC evolutionary biology 11, 1 (2011).

[57] P. Garcia-Galindo and S. E. Ahnert, Phenotypic plasticity can be an evolutionary response to fluctuating environments, bioRxiv, 2024.10.02.614758 (2024).

[58] M. Schmutzer, P. Dasmeh, and A. Wagner, Frustration can limit the adaptation of promiscuous enzymes through gene duplication and specialisation, Journal of Molecular Evolution 92, 104 (2024).

[59] J. L. Payne, F. Khalid, and A. Wagner, RNA-mediated gene regulation is less evolvable than transcriptional regulation, PNAS 115, E3481 (2018).

[60] J. Aguilar-Rodríguez, J. L. Payne, and A. Wagner, A thousand empirical adaptive landscapes and their navigability, Nat. Ecol. Evol. 1, 0045 (2017).

