## Supplementary text for "A simple model captures key characteristics of biological non-deterministic genotype-phenotype maps"

Nora S. Martin

**S5 Threshold-based framework for ND GP maps** **17**

#### S1 Theory: robustness and evolvability in ND GP maps

##### S1.1 Implications of genetic correlations in ND GP maps

A GP map has 'genetic correlations' if it has local correlations in genotype space: in the deterministic case, this means that for a genotype mapping to $p$, its mutational neighbours are more likely to map to $p$ than an arbitrary genotype $h$ [1]. Since the probability that mutational neighbours conserve $p$ is given by robustness $\rho_p$, and the probability that an arbitrary genotype gives $p$ is given by the frequency $f_p$, genetic correlations can be detected by analysing if the robustness $\rho_p$ of a phenotype $p$ exceeds its frequency $f_p$ [1].

In ND GP maps, 'genetic correlations' have been defined by analogy: a GP map has genetic correlations if $\tilde{\rho}_p > \tilde{f}_p$ for all/most phenotypes $p$ [2, 3]. This can similarly be interpreted in terms of 'clustering' of $p$ in genotype space: the following argument demonstrates that $\tilde{\rho}_p > \tilde{f}_p$ means that there is a positive covariance between $P(p|g)$ and $P(p|g')$, where $g'$ is a mutational neighbour of $g$. To show this, let us begin by using the definition of $\tilde{\rho}_p$, and writing $\tilde{\rho}_p > \tilde{f}_p$, as:

$$\frac{1}{K^L \cdot \tilde{f}_p \cdot (K-1)L} \sum_g P(p|g) \sum_{g' \in \mathcal{N}_g} P(p|g') > \tilde{f}_p$$

Rearranging gives:

$$\frac{1}{K^L \cdot (K-1)L} \sum_{g,g' \in \mathcal{N}_g} P(p|g)P(p|g') > \tilde{f}_p^2$$

The expression on the LHS can be recognised as the mean of $P(p|g)P(p|g')$, averaged over all pairs of a genotype $g$ and its mutational neighbour $g'$. Similarly, on the RHS, $\tilde{f}_p$ is the average of $P(p|g)$ over all genotypes $g$ (or, equivalently over all mutational neighbours of all genotypes). This gives:

$$E[P(p|g)P(p|g')] > E[P(p|g)]E[P(p|g')]$$

And thus, a positive covariance between $P(p|g)$ and $P(p|g')$:

$$cov(P(p|g), P(p|g')) = E[P(p|g)P(p|g')] - E[P(p|g)]E[P(p|g')] > 0$$

Therefore, genetic correlations can be interpreted as a type of 'local clustering' in genotype space: if a genotype $g$ has a high/low $P(p|g)$ (relative to the mean prevalence of $p$, which is given by its frequency $\tilde{f}_p$), then its mutational neighbours $g'$ are more likely to also have high/low $P(p|g')$ than in a completely uncorrelated GP map.

Interestingly, this perspective on genetic correlations in ND GP maps highlights that 'genetic correlations' are related to Ancel & Fontana's [4] concept of plastogenetic congruence (as noted before in ref [2]). Plastogenetic congruence can be defined in several ways, for example as follows: let us consider a genotype $g$, for which phenotype $p$ is the highest-$P(p|g)$ phenotype in the ensemble. Then, in a GP map with plastogenetic congruence, $g$'s mutational neighbours $g'$ are likely to retain $p$ as a high-$P(p|g)$ phenotype, even if not as the *highest*-$P(p|g)$ phenotype. While this definition differs from that of genetic correlations, for example with its focus on the *highest*-$P(p|g)$ phenotype for a given genotype, it is clearly related.

#### S1.2 Limit on robustness and evolvability on the genotypic level

To put limits on genotypic robustness and evolvability in ND GP maps, let us exploit parallels with simpler GP maps without ND. In GP maps without ND, there is a simple trade-off between robustness and evolvability on the genotypic level [5]: each genotype has $(K-1)L$ neighbours, where $K$ is the alphabet size and $L$ the sequence length. Each neighbour can either be neutral and contribute $\Delta\rho_g = 1/((K-1)L)$ to the genotypic robustness, or be non-neutral. If the neighbour is non-neutral and produces a phenotype that is not already in the neighbourhood, the mutation can contribute $\Delta e_g = 1$ to the evolvability, otherwise the mutation contributes to neither robustness nor evolvability. Thus, the following bound applies to GP maps without ND [5]:

$$e_g + ((K-1)L)\rho_g \leq (K-1)L$$
$$\frac{e_g}{(K-1)L} + \rho_g \leq 1 \quad (1)$$

To apply this result to GP maps *with* ND, let us use parallels between genotypic robustness and evolvability in GP maps *with* and *without* ND: in a GP map with ND, each genotype maps to a distribution of phenotypes. Let us now construct a GP map without ND, by simply drawing a single phenotype from each genotype's ensemble and recording it. If a large number of such 'frozen' GP maps without ND is drawn from a GP map with ND, then the average robustness of a genotype over the set of 'frozen maps' $\langle\rho_g\rangle_{\text{frozen}}$ approximates its robustness in the full ND GP map $\tilde{\rho}_g$ [2]. The same approximate equivalence holds for genotypic evolvability. This equivalence holds by design, thereby ensuring consistency between GP maps with and without ND.

The approximate equivalence between ND GP maps and averages over specific GP maps without ND can now be applied to eq. 1: in each of the frozen maps without ND, the genotypic robustness $\rho_g$ and evolvability $e_g$ must satisfy $\frac{e_g}{(K-1)L} + \rho_g \leq 1$. Thus, the same inequality must hold for their mean over frozen maps:

$$\langle\frac{e_g}{(K-1)L} + \rho_g\rangle_{\text{frozen}} \leq 1$$
$$\frac{\langle e_g\rangle_{\text{frozen}}}{(K-1)L} + \langle\rho_g\rangle_{\text{frozen}} \leq 1$$

Then the approximate equivalence of the means over frozen maps ($\langle e_g \rangle_{\text{frozen}}$ and $\langle \rho_g \rangle_{\text{frozen}}$) with the ND GP map quantities ($\tilde{e}_g$ and $\tilde{\rho}_g$) can be applied to write:

$$\frac{\tilde{e}_g}{(K-1)L} + \tilde{\rho}_g \leq 1$$

Thus, genotypic robustness and evolvability in the ND GP map approximately satisfy the same trade-off as in the well-known case without ND.

#### S2 Further analyses of the three biophysical ND GP maps

##### S2.1 Phenotypic frequency distribution

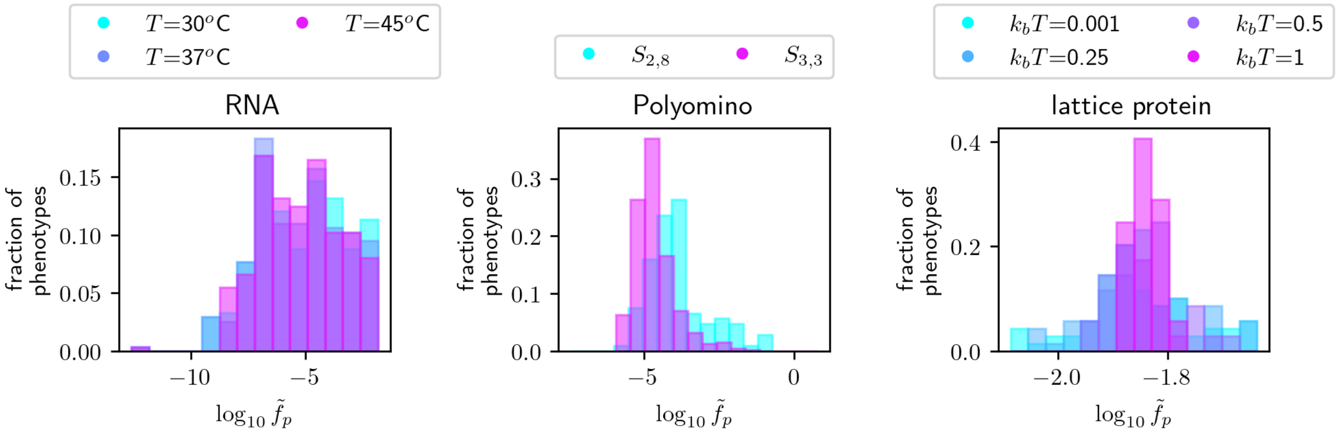

Figure S1: **Distribution of phenotypic frequencies in the three biophysical GP maps:** RNA secondary structure, Polyomino self-assembly and HP lattice proteins.

Fig S1 shows the distribution of phenotypic frequencies $\tilde{f}_p$ in the three biophysical models. All maps show phenotypic bias since different phenotypes have different phenotypic frequencies $\tilde{f}_p$. However, the details of the distribution differ in the three maps. This diversity in frequency distributions is consistent with work on GP maps without ND, which has found a log-binomial [6] or log-normal [7] distribution for the case of RNA, but a distribution dominated by the lowest-frequency class in the GP map of Richard Dawkins' biomorphs [8].

##### S2.2 Genetic correlations - further data

As shown in section S1.1, there are two ways of detecting genetic correlations: either by testing whether the phenotypic frequency $\tilde{f}_p$ of a phenotype $p$ is higher than its robustness $\tilde{\rho}_p$, or testing for covariance between the probabilities $P(p|g)$ and $P(p|g')$ for pairs of mutational neighbours $g$ and $g'$. This section provides additional data on both options.

First, Fig. S2 shows the ratio between each phenotype's robustness $\tilde{\rho}_p$ and its frequency $\tilde{f}_p$ on a logarithmic scale. Since genetic correlations are present whenever $\tilde{\rho}_p/\tilde{f}_p > 1$, they can easily be detected by applying the criterion $\log_{10}(\tilde{\rho}_p/\tilde{f}_p) > 0$ (black line in the histograms). The phenotypes in the biophysical ND GP maps meet this criterion, with the exception of 303 zero-robustness phenotypes in the $S_{3,3}$ version of the Polyomino model, which all have low frequencies of $\tilde{f}_p < 3.4 \times 10^{-4}$. These exceptions aside, many phenotypes have $\log_{10}(\tilde{\rho}_p/\tilde{f}_p) \gtrapprox 1$, indicating that robustness values are $\gtrapprox 10$ times as high as the corresponding frequency values.

Secondly, let us turn to the alternative way of testing for genetic correlations: there is covariance between $P(p|g)$ and $P(p|g')$, where $g$ and $g'$ are mutational neighbours. This can be detected using Pearson correlation coefficients as a normalised measure of covariance. As expected, the correlation between $P(p|g)$ and $P(p|g')$

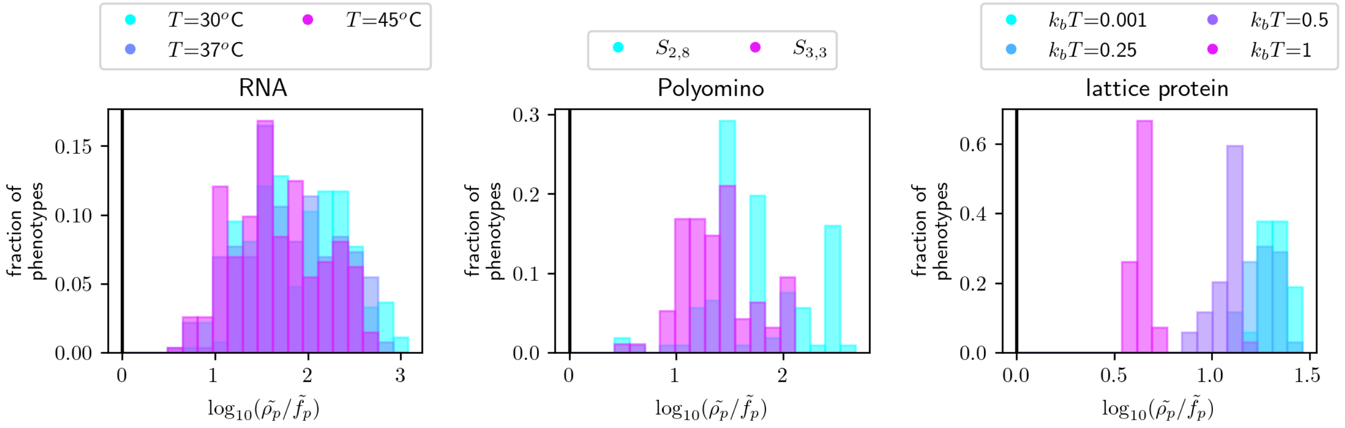

Figure S2: **Ratio between phenotypic robustness $\tilde{\rho}_p$ and frequency $\tilde{f}_p$ as an alternative method of quantifying genetic correlations in the biophysical GP maps:** the histograms show $\log(\tilde{\rho}_p/\tilde{f}_p)$. Since a GP map has genetic correlations if $\log(\tilde{\rho}_p/\tilde{f}_p) > 0$ (see black line at zero), this analysis confirms the conclusions from the main text: genetic correlations are present in all three maps, RNA, Polyomino and lattice protein. The only exception are zero-robustness phenotypes in the Polyomino model (303 in the $S_{3,3}$ case), which are not shown due to the logarithmic scale and all have low frequencies of $\tilde{f}_p < 3.4 \times 10^{-4}$.

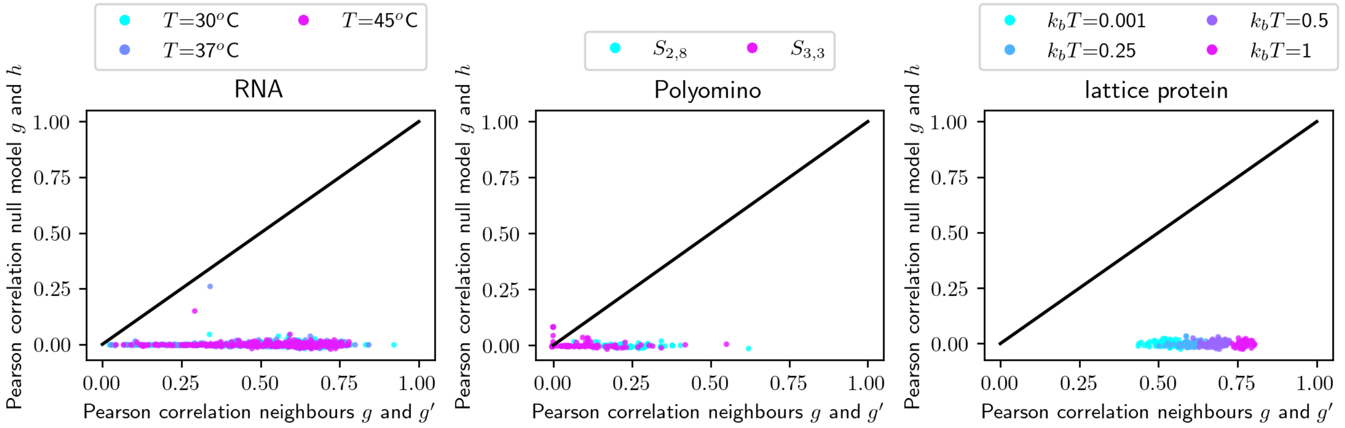

Figure S3: **Correlation between $P(p|g)$ and $P(p|g')$ for two mutational neighbours $g$ and $g'$ as a second method of detecting genetic correlations in the biophysical GP maps:** the plotted data were generated as follows: $10^4$ $g/g'$ pairs were generated at random. $P(p|g)$ and $P(p|g')$ were recorded for each pair and each phenotype. Then, for each phenotype $p$, a Pearson correlation coefficient was computed between the $P(p|g)$ data and the corresponding $P(p|g')$ data (x-axis). The zero-correlation null model (y-axis) was computed in the same way, except that a random genotype $h$ replaced the mutational neighbour $g'$. Since most correlation coefficients are positive and exceed the null model, the GP maps have genetic correlations. The only exceptions are low-frequency phenotypes in the Polyomino model (see text).

in the biophysical ND GP maps (Fig S3) tends to be higher when computed for two mutational neighbours $g$ and $g'$ than for two arbitrary genotypes $g$ and $h$. Exceptions, where the correlation is negative, lower than the null model or cannot be computed because $P(p|g) = 0$ for all sampled genotypes, exist for the low-frequency phenotypes in the Polyomino model: 18 phenotypes in the $S_{2,8}$ map (with $\tilde{f}_p < 8.2 \times 10^{-5}$) and 325 phenotypes in the $S_{3,3}$ map (with $\tilde{f}_p < 3.4 \times 10^{-4}$), including all 303 zero-robustness phenotypes.

##### S2.3 Applying recent theory to the frequency-robustness data

This section applies recent theoretical approximations to the phenotypic-robustness-frequency data in the three biophysical ND GP maps: First, Sappington & Mohanty [3] propose an upper bound for the phenotypic robustness $\tilde{\rho}_p$ of phenotypes of a given frequency $\tilde{f}_p$. This upper bound is plotted together with the simulation data in the upper row of Fig. S4. The proposed upper bound is consistent with the data, despite the

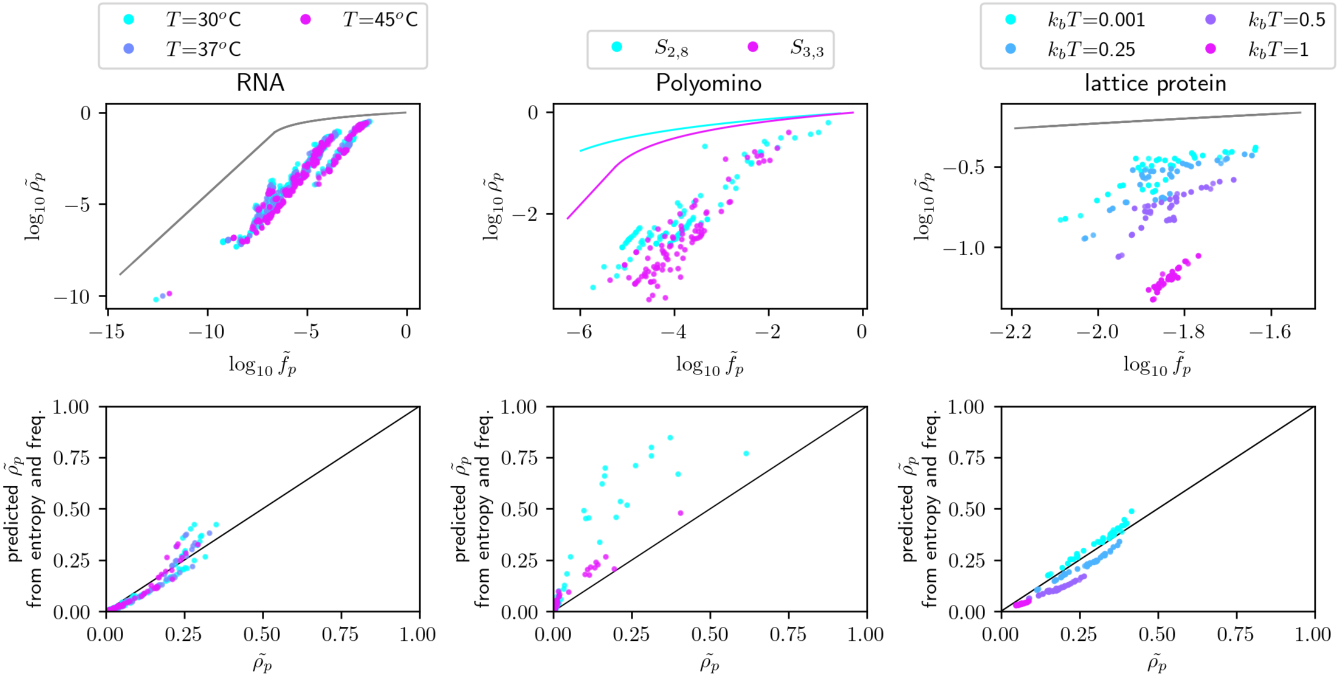

Figure S4: **Sappington & Mohanty's [3] theory on phenotypic robustness applied to the three biophysical ND GP maps: (first row)** Logarithmic plot of phenotypic robustness $\tilde{\rho}_p$ against phenotypic frequency $\tilde{f}_p$ (scatter points), together with Sappington & Mohanty's [3] upper bound (drawn as a line). Note that the two versions of the Polyomino model have different upper bounds since they differ in their alphabet size $K$ and sequence length $L$. **(second row)** Sappington & Mohanty's [3] approximate relationship between phenotypic robustness $\tilde{\rho}_p$, phenotypic frequency $\tilde{f}_p$ and phenotypic entropy $\tilde{S}_p$ is tested: the predicted phenotypic robustness (RHS of eq. 2) is plotted against the computational phenotypic robustness $\tilde{\rho}_p$. The black line illustrates $x = y$, i.e. a perfect prediction.

approximations made in its derivation (esp. a binary assumption about phenotypic ensemble frequencies: that each phenotype either has a fixed ensemble frequency $x$ or does not exist in an ensemble). Secondly, Sappington & Mohanty [3] derive a second approximation: that the phenotypic robustness $\tilde{\rho}_p$ depends on the phenotypic frequency $\tilde{f}_p$, as well as on the entropy of a phenotype $\tilde{S}_p$, as [3]:

$$\tilde{\rho}_p \approx \frac{K^L \tilde{f}_p \tilde{S}_p \exp(-\tilde{S}_p)}{L \log(K)} \tag{2}$$

Here, the entropy $\tilde{S}_p$ quantifies whether the probability weight of a phenotype $p$ is concentrated on a small or large number of genotypes and is defined as [3]:

$$\tilde{S}_p = - \sum_{\text{genotypes } g} \frac{P(p|g)}{\tilde{f}_p K^L} \log \frac{P(p|g)}{\tilde{f}_p K^L} \tag{3}$$

To test the relationship from eq. 2, the predicted phenotypic robustness (RHS of eq. 2) is plotted against the true phenotypic robustness $\tilde{\rho}_p$ (second row of Fig. S4): In agreement with Sappington & Mohanty's [3] data esp. on RNA, the predicted phenotypic robustness is highly correlated with robustness.

##### S2.4 Distance between the 'peaks' of different phenotypes' probabilities

Let us analyse one feature that only exists in *ND* GP maps: the genotypes that maximise $P(p|g)$ for different phenotypes $p$. In the ND case, genotypes mapping to $p$ do so with different probabilities $P(p|g)$. Thus, there is a 'peak' for each phenotype $p$: a genotype $g$ that maximises $P(p|g)$ for that $p$. This raises the question of how the peak genotypes of different phenotypes are spread in genotype space. To illustrate this, Fig S5 shows the Hamming distance between pairs of genotypes that maximise $P(p|g)$ for two distinct phenotypes $p$ and $q$. The histograms show that these 'peaks' almost have the same Hamming distance as random genotypes

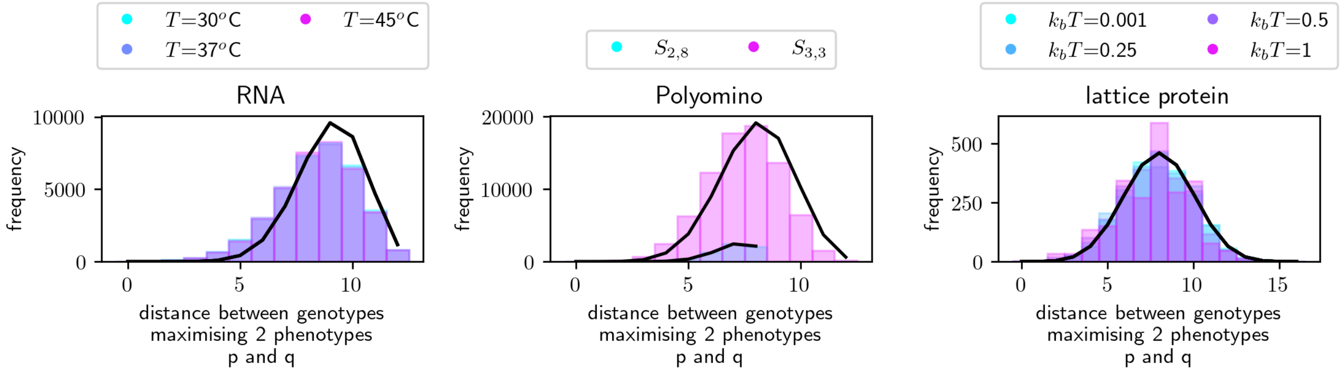

Figure S5: **Distance between 'peaks'**: for each of the biophysical GP maps, the genotypes that maximise $P(p|g)$ for one phenotype $p$ were identified (to avoid rounding error artefacts, the analysis was restricted to phenotypes $p$ with at least one genotype $g$ with $P(p|g) > 10^{-6}$). If there were several genotypes with $P(p|g) > \max_g(P(p|g)) - 10^{-7}$, one of them was selected at random; for the Polyomino model, where several genotypes can define identical assembly graphs and thus identical probability distributions, the assembly graph maximising $P(p|g)$ for a given $p$ was selected, and then a random genotype mapping to this assembly graph was chosen. Once one 'peak' genotype per phenotype was selected, the Hamming distance between pairs of such 'peak' genotypes was computed and is shown here. For context, the black line illustrates the Hamming distance distribution between two arbitrary genotypes.

(see lines in Fig S5), indicating that they are not highly clustered in genotype space. Future work could investigate this in more detail and analyse the evolutionary implications of these peak-peak distances: It is likely that evolutionary transitions are faster between two phenotypes $p$ and $q$ if their peaks are closer.

This analysis is closely related to work on prototype sequences in the lattice protein model. These prototype sequences are defined based on their robustness in the deterministic GP map but often have the highest stability for a given structure [9] and would thus correspond to the 'peaks' in the ND GP map. These prototype sequences are a little less diverse than random sequences [10], consistent with the results in Fig. S5.

#### S2.5 ND GP map characterisation for different ensemble cut-offs in the Polyomino model

In the Polyomino model, a cut-off is used when processing the self-assembly simulations: the self-assembly process was repeated 500 times and thus the ensemble frequency was estimated to be $P(p|g) = x/500$ for a structure $p$ appearing $x$ times. However, since reliable estimates cannot be made for structures appearing very rarely, a cut-off is applied and structures with less than $x = 5$ appearances are treated as 'undefined' phenotypes (see methods section in the main text). To investigate the impact this cut-off may have on the ND GP map analysis, the analysis in Fig. 3 of the main text is repeated for two different cut-offs, $x = 3$ and $x = 10$ (shown in Fig. S6, with the $x = 5$ data also shown for comparison). On a qualitative level, the conclusions are unaffected.

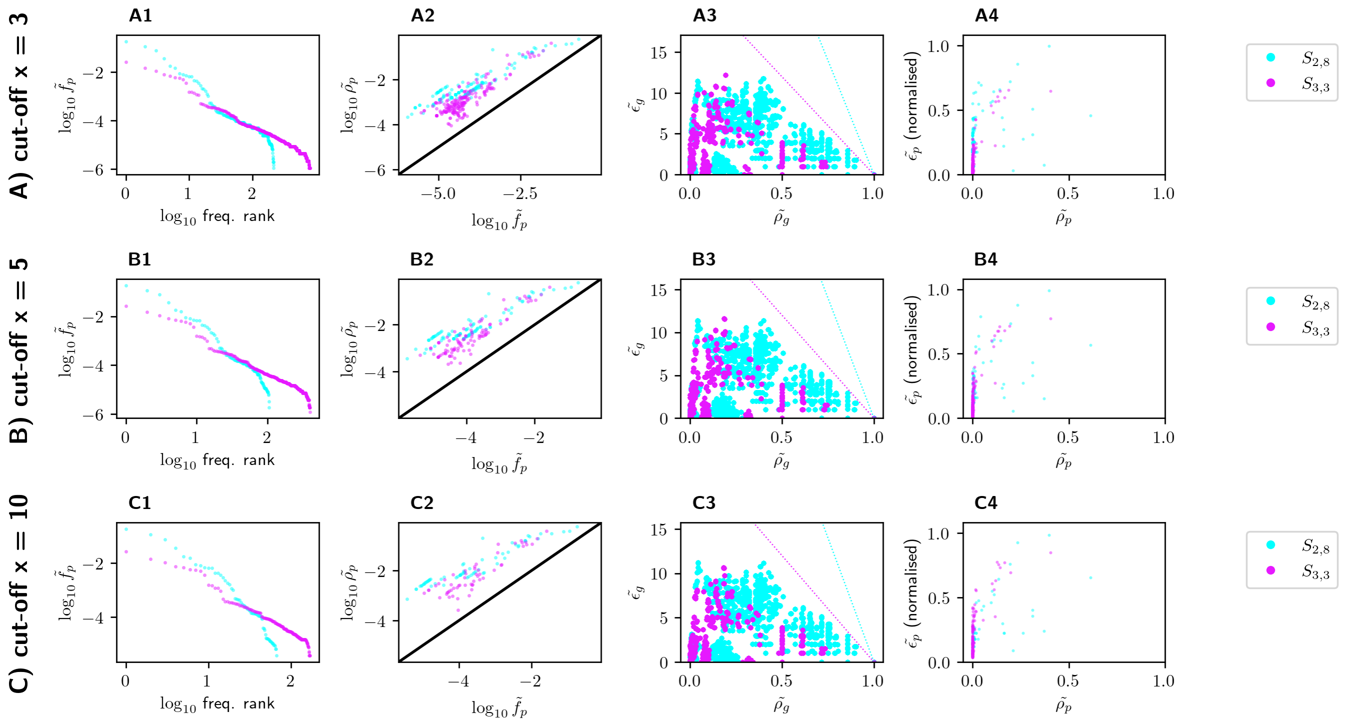

Figure S6: **Polyomino ND GP map analysis for different thresholds $x$ in the ensemble calculation:** The ensemble frequencies in the Polyomino model are based on the assembled structures found after 500 simulated assembly processes per assembly graph. Since the frequencies of low-$P(p|g)$ structures are difficult to estimate reliably, assemblies appearing fewer than $x$ times were treated as 'undefined'. To analyse the impact of this choice, this figure repeats the ND GP map analysis for different choices of $x$, one in each row: **(A)** x = 3, **(B)** x = 5 (as in the main text, shown for context), **(C)** x = 10.

### S3 Further analysis of the synthetic GP map

#### S3.1 Applying recent theory to the frequency-robustness data

As before for the biophysical maps (section S2.3), let us apply recent theoretical approximations for the phenotypic-robustness-frequency relationship to the synthetic ND GP maps: First, the phenotypic-robustness-frequency data is shown together with Sappington & Mohanty's [3] upper bound (column 1 of Fig. S7). The approximate upper bound is consistent with the computational data for the synthetic ND GP map, and for low stochasticity $T$, i.e. close to the deterministic limit, the computational data lies close to this upper bound.

Secondly, let us turn to Sappington & Mohanty's [3] approximate robustness predictions in terms of frequency $\tilde{f}_p$ and entropy $\tilde{S}_p$ (eq. 2). In column 2 of Fig. S7, this prediction is plotted against the computational robustness data for the synthetic ND GP map. As before for the biophysical ND GP map, this approximation works well. However, it cannot fully account for the impact stochasticity $T$ has on robustness. This may be because the derivation makes specific assumptions about the ND GP map (for example, that $P(p|g)$ only takes one non-zero value for each $p$). How well this approximation works may depend on the typical ensemble properties of the GP map, which in turn are affected by stochasticity $T$: for example, the typical entropy of a phenotype $\tilde{S}_p$ increases markedly with stochasticity $T$ (see column 3 of Fig. S7), as noted before for high-temperature versions of the RNA or spin-glass ND GP maps [3].

#### S3.2 High-stochasticity limit $T \to \infty$

In the high-stochasticity limit $T \to \infty$, any genotype $g$ maps to any phenotype $p$ with the same ensemble probability $P(p|g) = 1/n_p$. This is so simple that ND GP map characteristics can be computed analytically: Since there are $K^L$ genotypes in the ND GP map, the phenotypic frequency can be written as:

$$\tilde{f}_p = \frac{1}{K^L} \sum_g \frac{1}{n_p} = \frac{1}{n_p}$$

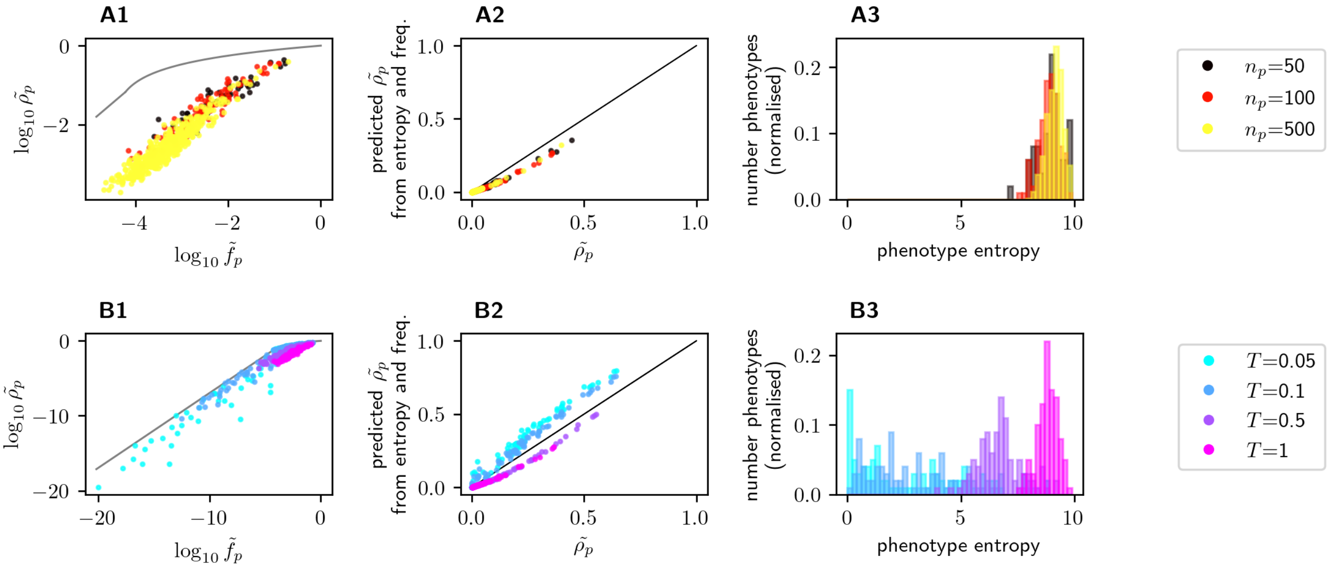

Figure S7: **Sappington & Mohanty's [3] theory on phenotypic robustness applied to the synthetic ND GP maps: (column 1)** Logarithmic plot of phenotypic robustness $\tilde{\rho}_p$ against phenotypic frequency $\tilde{f}_p$ (scatter points), together with Sappington & Mohanty's [3] upper bound (drawn in grey). **(column 2)** Sappington & Mohanty's [3] approximate relationship between phenotypic robustness $\tilde{\rho}_p$, phenotypic frequency $\tilde{f}_p$ and phenotypic entropy $\tilde{S}_p$ is tested by plotting the predicted phenotypic robustness (RHS of eq. 2) against the computational phenotypic robustness $\tilde{\rho}_p$. The black line illustrates $x = y$, i.e. a perfect prediction. **(column 3)** The entropy (eq. 3) is computed for each phenotype and shown as a histogram, illustrating that the entropy increases with $T$. This observation is consistent with the intuition that phenotypes spread out further in genotype space in the highly non-deterministic limit. **(rows)** As in Fig. 5 in the main text, row A shows ND GP maps with varying $n_p$ (and $T$ set to the mean difference in $G_{p,\vec{g}}$ between the two highest-probability phenotypes), and row B shows ND GP maps with fixed $n_p = 100$ and varying $T$.

Since there are $(K-1)L$ mutational neighbours per genotype, the phenotypic robustness can be written as:

$$\tilde{\rho}_g = \frac{1}{(K-1)L} \sum_p \frac{1}{n_p} \sum_{g' \in \mathcal{N}_g} \frac{1}{n_p} = \frac{1}{n_p}$$

The genotypic robustness is given by:

$$\tilde{\rho}_p = \frac{1}{K^L \cdot \tilde{f}_p \cdot (K-1)L} \sum_g \frac{1}{n_p} \sum_{g' \in \mathcal{N}_g} \frac{1}{n_p} = \frac{1}{n_p}$$

The genotypic evolvability (assuming $L \gg 1$) is given by:

$$\tilde{\epsilon}_g = \sum_p \frac{1}{n_p} \sum_{p' \neq p} \big(1 - \Pi_{g' \in \mathcal{N}_g} (1 - \frac{1}{n_p})\big) = (n_p - 1)\big(1 - (1 - \frac{1}{n_p})^{(K-1)L}\big) \approx (n_p - 1)$$

Finally, the phenotypic evolvability (assuming $L \gg 1$) is given by:

$$\tilde{\epsilon}_p = \sum_{p' \neq p} \big(1 - \Pi_g \Pi_{g' \in \mathcal{N}_g} (1 - \frac{1}{n_p}\frac{1}{n_p})\big) = (n_p - 1)\big(1 - (1 - \frac{1}{n_p^2})^{K^L(K-1)L}\big) \approx (n_p - 1)$$

#### S3.3 Deterministic GP map

The synthetic model, in addition to being a simple ND GP map, also defines a GP map without ND in the deterministic limit $T \to 0$. In this deterministic GP map, each genotype $\vec{g}$ maps to the phenotype $p$ with lowest $G_{p,\vec{g}}$; in the exceptional case that the two lowest-$G_{p,\vec{g}}$ phenotypes differ by less than $10^{-4}$, the genotype is treated as 'undefined', in line with conventions for ties in the HP model [1]. Thus, for a given

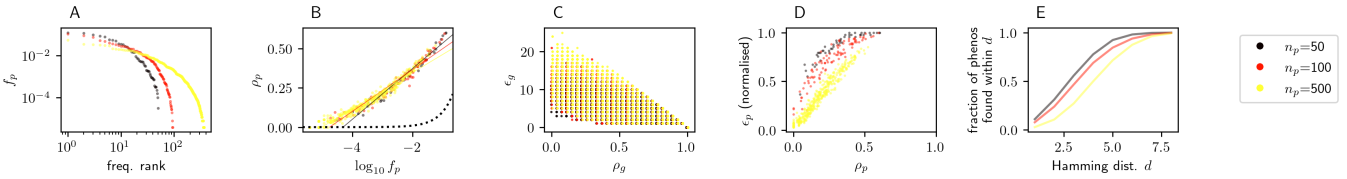

Figure S8: **The synthetic GP map in its deterministic limit ($T = 0$) displays the 'universal' properties of GP maps without ND (see reviews [11, 12]): (A)** The relationship between phenotypic frequency $f_p$ and frequency rank shows clear phenotypic bias. **(B)** Phenotypic robustness $\rho_p$ is plotted against frequency $f_p$ on a lin-log scale, showing genetic correlations ($\rho_p > f_p$ for the majority of phenotypes, highlighted by the black dotted line showing $\rho_p = f_p$), and a good fit with a log-linear function (fitted lines). **(C)** There is a trade-off between genotypic evolvability $e_g$ and robustness $\rho_g$ (x-values are shifted by up to $0.1/((K-1)L)$) to show overlapping data points more clearly). **(D)** There is a positive correlation between phenotypic robustness $\rho_p$ and evolvability $e_p$ (here normalised by the maximum possible evolvability $m_p - 1$, where $m_p \leq n_p$ is the number of phenotypes that appear at least once in the deterministic GP map). **(E)** Shape-space covering (following convention of ref [13]): how many distinct phenotypes are present within a Hamming distance $d$ from a given initial genotype? Here, the mean over $10^2$ randomly selected initial genotypes is shown, normalised by the highest possible value of $m_p - 1$.

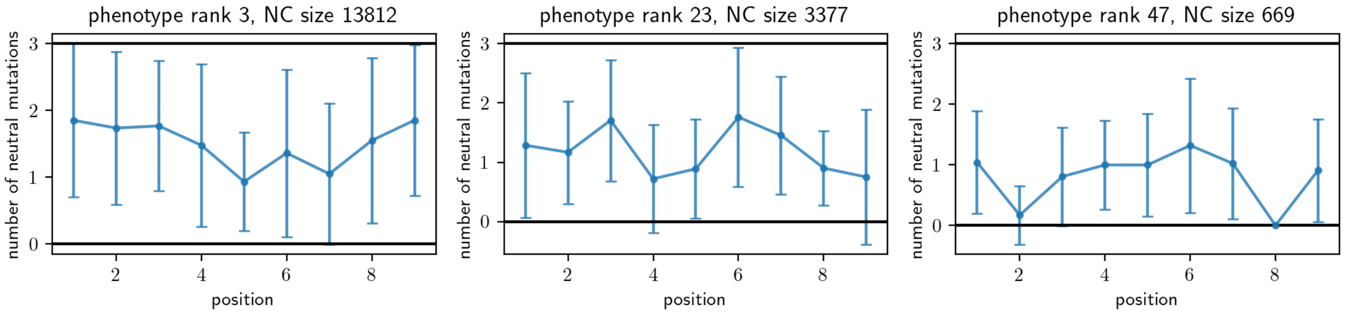

Figure S9: **Does the synthetic GP map in its deterministic limit ($T = 0$, here for $n_p = 100$) have position-specific sequence constraints (which are central in current minimal models of deterministic GP maps [14–16]):** position-specific sequence constraints are visualised for three phenotypes of different frequency ranks, one phenotype in each subplot (see titles). To quantify sequence constraints computationally, I first selected a random genotype $g$ mapping to $p$ and extracted the corresponding neutral component (NC), the set of genotypes mapping to $p$ that are accessible through neutral mutations from $g$. Then, following Weiß & Ahnert [17], I evaluated the number of neutral mutations at each sequence position position as one indicator of sequence constraints. For each position (x-axis), the mean and standard deviation over all genotypes in the NC are shown (y-axis). The plots show that different sequence positions in a NC have different sequence constraints, but there is considerable variability around this mean, even within a single NC.

sequence length $L$, only one free parameter remains: the number of phenotypes $n_p$. However, one change was made to the simple model introduced in the main text: the alphabet size was raised to $K = 4$ (with sequence length $L = 9$), to obtain a higher number of mutations per sequence position and thus a more nuanced picture of sequence constraints. In order to generalise the model to an arbitrary alphabet size $K$, the parameter vectors $\vec{v_p}$ need to be extended to length $(K-1)L$. Then, $G_{p,g}$ can be computed as:

$$G_{p,g} = \sum_{\text{non-zero seq. positions } i} (\vec{v_p})_{i+(\vec{g}_i - 1)L}$$

Fig. S9 analyses three versions of this GP map, for $n_p = 50$, $n_p = 100$ and $n_p = 500$, finding all central 'universal' features of deterministic GP maps (see ref [11]): first, there is strong phenotypic bias, since different phenotypes differ in their phenotypic frequencies $f_p$ by orders of magnitude (Fig. S9A). Secondly, the phenotypic frequency-robustness relationship is well-described by a log-linear fit (Fig. S9B) and the phenotypic robustness is higher than the frequency for the majority of phenotypes (exceptions: 3 out of the 94 phenotypes appearing at least once in the $n_p = 100$ case, 27 out of 368 in the $n_p = 500$ map; all of these phenotypes are rare with $f_p \leq 4/K^L$ and the majority has zero robustness simply because they are only generated by a single genotype). Thus the deterministic version of the synthetic map has genetic correlations. Thirdly, the relationship between *genotypic* evolvability and robustness shows a trade-off (Fig. S9C), but that of *phenotypic* evolvability and robustness is positive (Fig. S9D). Finally, the synthetic model satisfies the space-shape-covering property, i.e. most phenotypes are available within a comparatively small Hamming distance from an arbitrary genotype (Fig. S9E).

Thus, in the deterministic limit, the synthetic GP map displays the well-studied 'universal' characteristics. Simple models reproducing these shared characteristics have typically had variable sequence constraints as a key feature: each sequence position is characterised by a fixed capacity for phenotype-conserving mutations (the 'versatility' [7]), which only depends on the phenotype [7, 8, 14, 17, 18]. This prompts the question of whether the synthetic GP map similarly has variable sequence constraints as a defining feature. Fig S9 visualises these sequence constraints for three phenotypes. The sequence constraints, quantified by the mean number of neutral mutations, show some position-dependence, as in 'sequence-constraint-based' models, but there is considerable variation around the mean. Thus, a 'sequence-constraint-based' model, which mainly uses mean versatilities to characterise a NC, may not approximate this GP map and its structure well. However, some variation around the mean can be accounted for within the framework of 'sequence-constraint-based' models, for example when extrapolating from versatilities to NC sizes [18]. Small differences in versatility are also present in a toy model called the 'RNA-like GP map model' [15]. The question of when a 'sequence-constraint-based' model stops being a useful approximation needs further investigation.

If the deterministic version of the synthetic GP map does not correspond to a 'sequence-constraint-based' model, this would mean that 'sequence-constraint-based' models are just one of several ways of generating a GP map with the well-studied 'universal' characteristics of deterministic GP maps. In fact, the error bars in Fig S9 may be interesting themselves since they represent a type of epistasis: the number of neutral mutations at a particular site is highly genotype-dependent, even among genotypes in a single NC. This epistasis may be of biological relevance: some variability in the position-specific versatility is already present in the RNA GP map [18], even though this GP map has been approximated based on its sequence constraints [7, 18]. Moreover, empirical evidence [19] and computational models fitted to sequence families [20] suggest that mutations, which are only neutral for some genotypes in a sequence family, may be common.

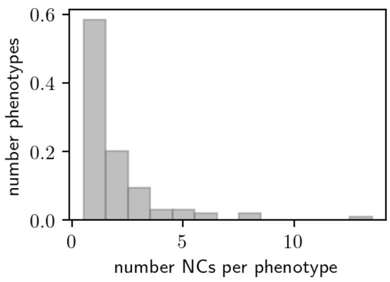

Figure S10: **Does the synthetic GP map in its deterministic limit ($T = 0$, here for $n_p = 100$) have NC fragmentation?:** the synthetic deterministic GP map has NC fragmentation, i.e. more than one mutationally connected NC per phenotype, for over 40% of all phenotypes.

Going beyond the scale of a NC, there is an additional feature that the synthetic GP map shares with biophysical models like RNA: the fact that the set of genotypes mapping to a single phenotype is made up of several mutationally disconnected subsets or NCs (Fig S10). This arises from another type of epistasis between neutral mutations [21] that is typically absent in the simple sequence-constraint-based toy models.

Thus, the synthetic model, by combining several additive functions with a non-linearity, produces a deterministic GP map with neutral epistasis. This observation is related to previous work on non-neutral mutations, showing that Boltzmann ensembles that depend on several additive scalars give rise to epistatic landscapes [22]. This feature of the synthetic model may be useful in the study of deterministic GP maps, offering a simple model that nevertheless contains neutral epistasis.

### S4 Variations of the synthetic ND GP map model

So far, the analysis has focused on one possible definition of a synthetic ND GP map. However, the non-linear function defining the map (Eq. 1 in the main text), the method of generating the phenotype-dependent parameters $\vec{v}_p$ and the alphabet size $K$ could have been chosen differently. In this section, the analysis is repeated with different options.

#### S4.1 ND GP maps based on different non-linear functions

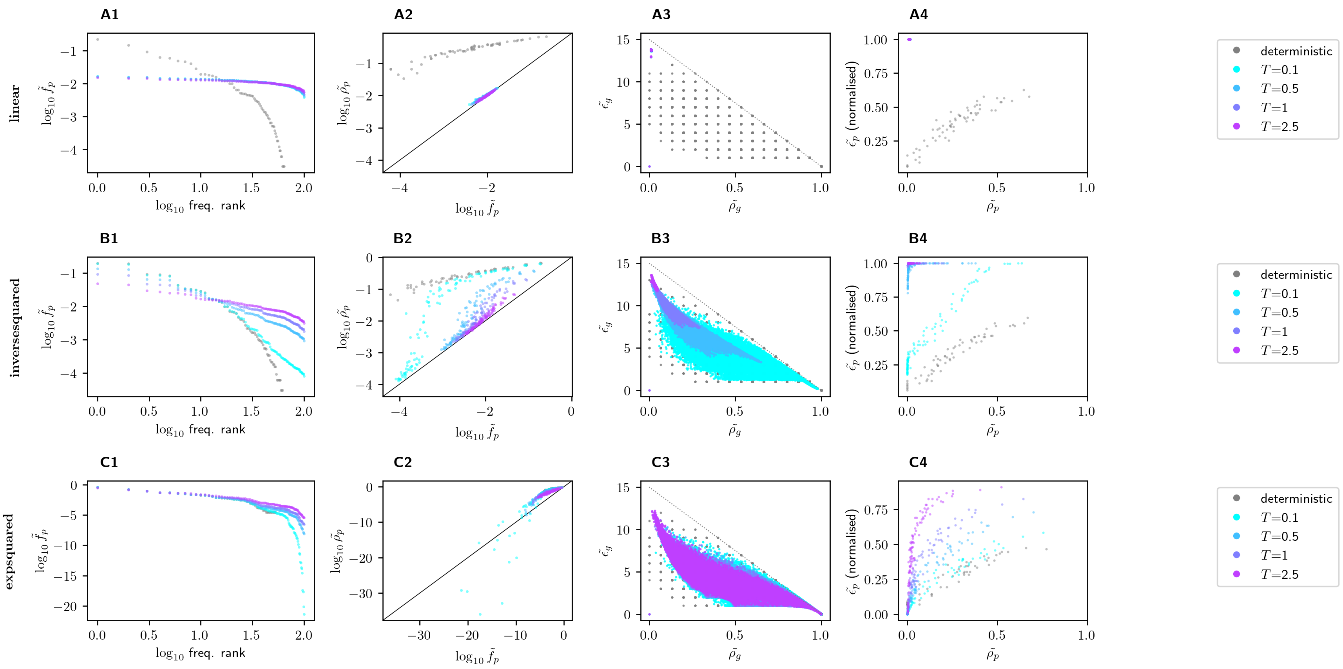

Figure S11: **ND GP map analysis for alternative versions of the synthetic ND GP map, based on different functional forms:** each row focuses on one alternative functional form in the ND GP map definition, which are described in more detail in the text (all with $n_p = 100$, and a range of values for the stochasticity $T$; the deterministic map is included in grey for context, even though this limit does not depend on the functional form): **(row A)** a 'linear' scaling with non-linearity in the y-intercept and normalisation only, **(row B)** an inverse-squared scaling as one example of a power law, **(row C)** a Gaussian (i.e. 'exponential-squared') scaling as one example that decays more quickly than the Boltzmann-like scaling used in the main text. As in Fig 5 in the main text, each column analyses one specific aspect of the GP map: **(1)** The phenotypic frequency $\tilde{f}_p$ is plotted against the frequency rank, showing phenotypic bias. This bias is much smaller in the linear map, spanning less than an order of magnitude, compared to all other examples, where the bias ranges across several orders of magnitude in the low-$T$ limit. **(2)** The phenotypic robustness $\log \tilde{\rho}_p$ is plotted against the phenotypic frequency $\log \tilde{f}_p$ ($x = y$ is drawn as a black line to guide the eye). Since $\log \tilde{\rho}_p > \log \tilde{f}_p$ for the majority of phenotypes, the ND GP maps have genetic correlations, but these correlations can be weak, for example in the linear map. **(3)** The genotypic evolvability $\tilde{e}_g$ is plotted against the genotypic robustness $\tilde{\rho}_g$. There are no high-$\tilde{e}_g$-high-$\tilde{\rho}_g$ genotypes, consistent with the upper bound from section S1.2 (shown as a grey dotted line; the data is below the bound when accounting for numeric errors of up to $10^{-4}$). While both low-$\tilde{e}_g$-high-$\tilde{\rho}_g$ and high-$\tilde{e}_g$-low-$\tilde{\rho}_g$ genotypes exist in most maps, the linear map only contains high-$\tilde{e}_g$-low-$\tilde{\rho}_g$ genotypes (with the exception of the all-0 genotype, which is a low-evolvability-low-robustness genotype mapping to the undefined phenotype). **(4)** The normalised phenotypic evolvability $\tilde{e}_p$ is plotted against the phenotypic robustness $\tilde{\rho}_p$. This relationship is generally positive, except for saturation effects. This saturation is especially pronounced in the 'linear' case, where all phenotypes reach an evolvability within 0.002% of the maximum. The different characteristics of the 'linear' map can be explained by the fact that this map is closest to the trivial map with $P(p|g) \approx 1/n_p$, since the ensemble frequency calculations do not amplify the genotype-and phenotype-dependent differences in $G_{p,g}$ as much as the other functions.

First, let us change the functional form of the non-linear part of the synthetic GP map, which takes in one scalar $G_{p,\vec{g}}$ for each phenotype $p$ and translates this to an ensemble probability for each phenotype $p$ (Eq. 1 in the main text). Not all non-linear function can be chosen, since the final ensemble needs to be a valid probability distribution, with $P(p|\vec{g}) \geq 0$ and $\sum_p P(p|\vec{g}) = 1$. These requirements still leave many options, and I will focus on the following four functional forms:

**1) 'Linear with normalisation'**: a 'true' linear function of the form $P(p|g) = aG_{p,\vec{g}} + b$ would not satisfy the normalisation and non-negativity criteria of a probability distribution. However, a similar (albeit technically non-linear) function satisfies both requirements:

$$P(p|g) = \frac{G_{\max,\vec{g}} - G_{p,\vec{g}} + T}{\sum_q (G_{\max,\vec{g}} - G_{q,\vec{g}} + T)}$$

Here, to guarantee non-negativity of the probabilities $P(p|g)$, $G_{\max,\vec{g}}$ is the maximum value of $G_{q,\vec{g}}$ over all phenotypes $q$ for the given genotype $\vec{g}$. The parameter $T$ is the only free parameter since any additional pre-factor would cancel between the numerator and denominator. In two respects, this functional form mirrors that from the main text: first, high-$G_{q,\vec{g}}$ phenotypes have lower probabilities and, secondly, these probability differences become weaker with increasing $T$. This can be seen most clearly in the limit $T \to \infty$, giving $P(p|g) \to 1/n_p$ for all phenotypes $p$ as before (section S3.2). Despite these parallels with the 'Boltzmann-like' function in the main text, there are also differences: most importantly, the suppression of high-$G_{q,\vec{g}}$ is exponential in the 'Boltzmann-like' function, but only linear here.

**2) 'Inverse-squared'** : as a second option, let us consider a non-linear function that decreases more quickly with $G_{q,\vec{g}}$ than the linear one, but less quickly than the exponential one, thereby forming an intermediate. I chose a polynomial of order two with the following form:

$$P(p|g) = \frac{(G_{p,\vec{g}} - G_{\min,\vec{g}} + T)^{-2}}{\sum_q (G_{q,\vec{g}} - G_{\min,\vec{g}} + T)^{-2}} \tag{4}$$

Here, to get non-negative, finite probability values $P(p|g)$, $G_{\min,\vec{g}}$ is the minimum value of $G_{q,\vec{g}}$ over all phenotypes $q$ for the given genotype $\vec{g}$. As before, high-$G_{q,\vec{g}}$ phenotypes have lower probabilities and, secondly, these probability differences become weaker with increasing $T$.

**3) 'Gaussian'** : as a third alternative, let us take a functional form that decreases even more quickly with $G_{q,\vec{g}}$ than the exponential 'Boltzmann-like' one. I chose a Gaussian of the following form:

$$P(p|g) = \frac{\exp(-(G_{p,\vec{g}} - G_{\min,\vec{g}})^2/T)}{\sum_q \exp(-(G_{q,\vec{g}} - G_{\min,\vec{g}})^2/T)} \tag{5}$$

As before, high-$G_{q,\vec{g}}$ phenotypes have lower probabilities and these probability differences become weaker with increasing $T$. Note that the choice of non-linear function only becomes relevant for the ND GP map. In the deterministic GP map, the functional form does not matter and the GP map can be generated without knowing the function as follows: $P(p|g) = 1$ for the phenotype $p$ with the lowest $G_{p,\vec{g}}$ and $P(p|g) = 0$ for all other phenotypes.

Fig. S11 shows that these different maps all display the key shared features: First, some phenotypic frequencies $\tilde{f}_p$ are higher than others, i.e. there is phenotypic bias. Secondly, phenotypic robustness $\tilde{\rho}_p$ is typically higher than frequency $\tilde{f}_p$, i.e. the maps have genetic correlations. Thirdly, there are no high-robustness-high-evolvability *genotypes*, but *phenotypic* evolvability increases with phenotypic robustness (unless there is saturation). However, there are also prominent differences: most notably, the 'linear' scaling only gives weak phenotypic bias, weak genetic correlations and very low robustness. A more detailed analysis of the different models (Fig. S12) reveals that this is simply because the linear model does not produce strong bias even within a single genotype's ensemble. In other words, even for small $T$, it is already close to the high-$T$ limit in the other maps. Thus, among the different ND GP maps, it is most similar to the trivial GP map, in which each ensemble frequency is $P(p|g) = 1/n_p$ regardless of the phenotype $p$ and genotype $g$ (see section S3.2).

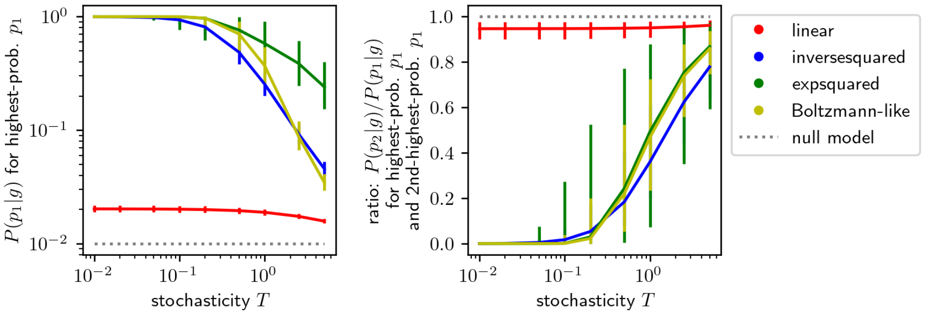

Figure S12: **Ensemble probabilities in versions of the synthetic ND GP map based on different functional forms:** this figure highlights the differences between the ensembles defined by the three non-linear functions in Fig. S12 as well as by the Boltzmann-inspired function from the main text (for $n_p = 100$). **(A)** For a range of $T$, the synthetic ND GP maps are built and the highest probability in the ensemble of each genotype $g$ is recorded (i.e. $P(p_1|g)$ where $p_1$ is the phenotype minimising $G_{p,g}$). The median and quartiles over all genotypes $g$, except the all-zero genotype as an outlier, are plotted. **(B)** For a range of $T$, the synthetic ND GP maps are built and the ratio of the highest two probabilities in the ensemble of each genotype $g$ is recorded. Again, the median and quartiles over all genotypes $g$, except the all-zero genotype as an outlier, are plotted. In both (A) & (B), a grey dotted line shows the expectation from a trivial null model, in which each ensemble frequency is $P(p|g) = 1/n_p$ regardless of the phenotype $p$ and genotype $g$ (thus, a genotype encodes no phenotypic information). This model is the high-$T$ limit of all models, but the ND GP map based on a linear function already approaches this null model at low $T$. This explains the lower phenotypic bias and genetic correlations in this map in Fig. S11.

#### S4.2 ND GP maps with parameters initialised from different probability distributions

This section modifies a second aspect of the synthetic ND GP map: the distribution from which the parameters in the linear part are chosen ($\vec{v}_p$ in Eq. 2 in the main text). Let us focus on three alternatives: an exponential, a log-normal and a uniform distribution (all with standard parameters: a scale of one in the exponential distribution, a mean of zero & standard deviation of one in the normal distribution underlying the log-normal distribution, and an interval between zero and one in the uniform distribution). Fig S13 shows that the key shared features continue to be present in these maps. However, there are also differences: for example, the value of $T$ at which saturation in the robustness-evolvability relationship sets in, differs from case to case. This is not surprising since the parameters $\vec{v}_p$ are changed and this affects the range of possible 'energy' values $G$ thus the ratio between 'energy' and $T$ at fixed $T$ (see eq. 1 in the main text). Moreover, in the 'log-normal' version, there is a higher number of low-frequency phenotypes, many of which do not satisfy the criterion for genetic correlations. Note, however, that many of these phenotypes are too rare to be relevant for any evolutionary questions: for example, phenotypes with $\tilde{f}_p < 10^{-10}$ must have $P(p|g) < 10^{-5}$ for all genotypes (since a phenotype with $P(p|g) = 10^{-5}$ for a single genotype has $\tilde{f}_p = 10^{-5}/2^15 \approx 3 \times 10^{-10}$).

An additional difference becomes apparent when repeating the analysis from section S2.4, identifying the 'peak' genotypes maximising the ensemble probability $P(p|\vec{g})$ for different phenotypes $p$ (Fig S15): the genotypes that maximise the probabilities of two different phenotypes $p$ and $q$ are more similar than two arbitrary genotypes, especially for some of the maps initialised from the exponential and log-normal distributions. The following hypothesis could explain this finding: if the vector elements are drawn from exponential and log-normal distributions, the vector elements of $\vec{v}_p$ can have extreme outliers - values that are much higher than the mean. Then, if $\vec{v}_p$ for some phenotype $p$ has an extremely high value at position $i$, any genotype with a 1 at the $i^{th}$ position will have an extremely high probability for phenotype $p$, and a very low probability for any other phenotype $q$. Thus, the genotypes that maximise any phenotype except $p$ are all likely to have a zero at position $i$, which makes them more similar to each other than arbitrary genotypes. This hypothesis is consistent with the tests in Fig S15: in the exponential and the log-normal map, genotypes that maximise a phenotype $q$ have a more biased composition towards zero, and this bias is strongest at sites $i$, for which one phenotype $p$ has an extremely high vector element $(\vec{v}_p)_i$.

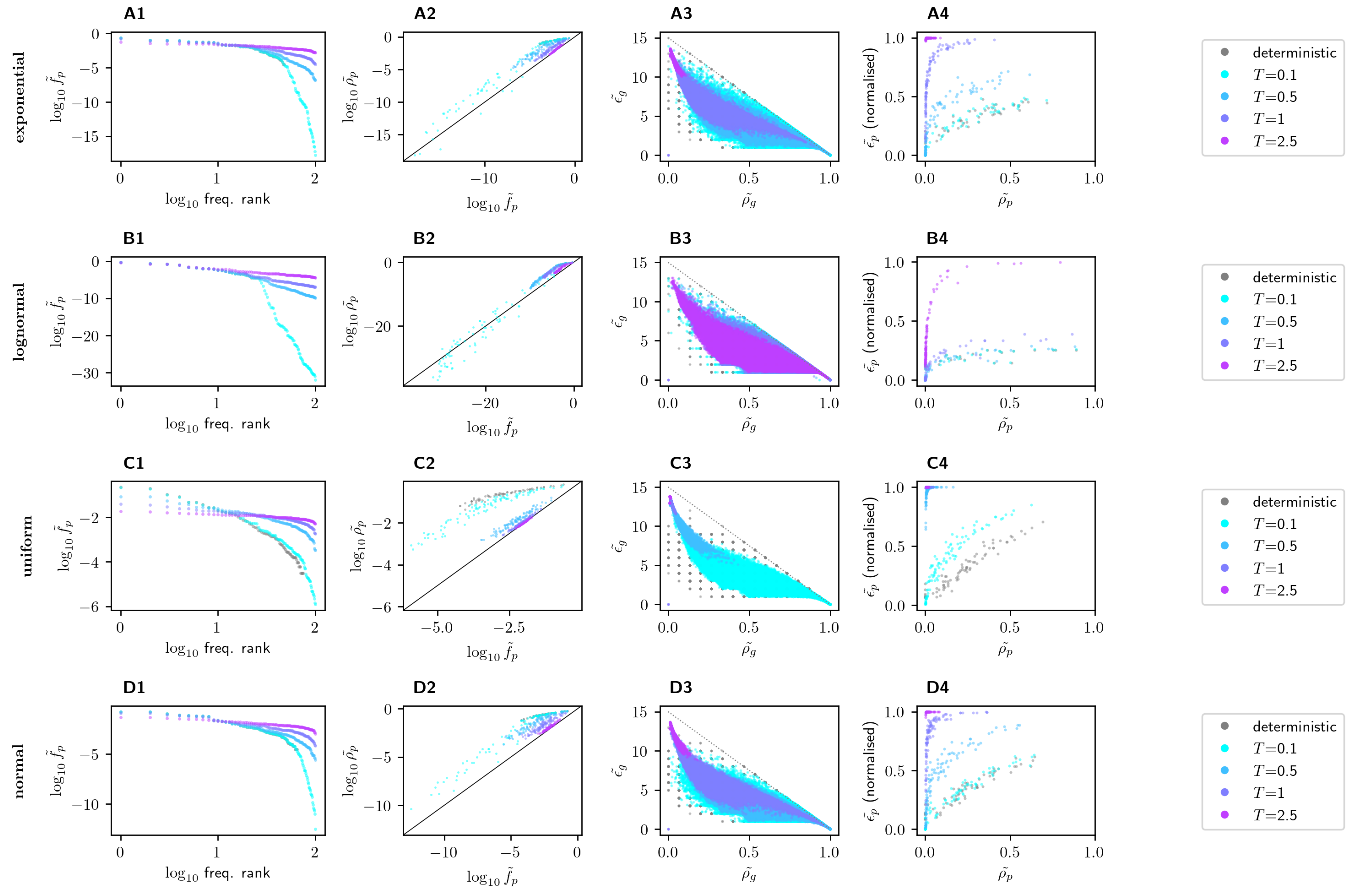

Figure S13: **ND GP map analysis of alternative versions of the synthetic ND GP map initialised from different probability distributions:** each row uses one alternative probability distributions to initialise the $\vec{v_p}$ parameters, as described in the text (all with $n_p = 100$, and a range of values for the stochasticity $T$; the deterministic limit, where each genotype maps to the lowest-$G$ structure, is shown in grey for comparison): **(row A)** exponential distribution, **(row B)** log-normal distribution, **(row C)** uniform distribution, **(row D)** normal distribution (as in main text, included for comparison). As in Fig 5 in the main text, each column analyses one specific aspect of the ND GP map: **(1)** The phenotypic frequency $\tilde{f}_p$ is plotted against the frequency rank, showing phenotypic bias. **(2)** The phenotypic robustness $\log \tilde{\rho}_p$ is plotted against the phenotypic frequency $\log \tilde{f}_p$ ($x = y$ is drawn as a black line to guide the eye). Since $\tilde{\rho}_p > \tilde{f}_p$ for the majority of phenotypes, the maps have genetic correlations. However, low-$T$ map initialised with a log-normal distribution has a high number of 42 exceptions, all exceptionally low-frequency phenotypes with $\tilde{f}_p < 3 \times 10^{-7}$. **(3)** The genotypic evolvability $\tilde{e}_g$ is plotted against the genotypic robustness $\tilde{\rho}_g$. There are no high-$\tilde{e}_g$-high-$\tilde{\rho}_g$ genotypes, consistent with the upper bound from section S1.2 (grey dotted line; the data is below the bound when accounting for numeric errors of up to $10^{-4}$). **(4)** The phenotypic evolvability $\tilde{e}_p$ is plotted against the phenotypic robustness $\tilde{\rho}_p$. Their relationship is generally positive, except for saturation effects, especially in the high-$T$ limit.

#### S4.3 ND GP maps with different alphabet sizes

As shown in section S3.3, the synthetic ND GP map can easily be extended to larger alphabet sizes $K$. Fig S16 shows two options: $K = 2$ and $K = 4$. Again, the key shared features are present, regardless of alphabet size.

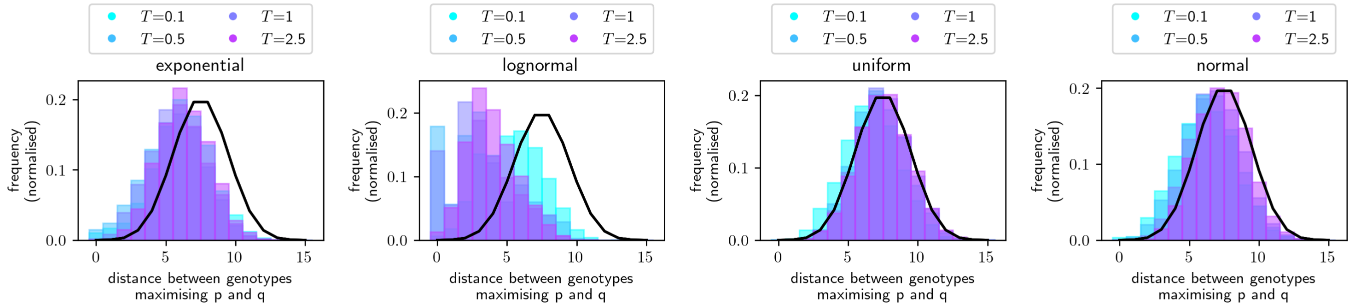

Figure S14: **Distance between 'peaks', i.e. genotypes that maximise $P(p|g)$ for one phenotype $p$**: For each of the GP maps defined in Fig. S13, the genotypes that maximise $P(p|g)$ for one phenotype $p$ are identified with the same conventions as in Fig. S5. The Hamming distance distribution between pairs of such 'peak' genotypes is shown as a histogram. The distribution for two arbitrary genotypes is shown in black for context. The histograms show that the 'peak' genotypes for two different phenotypes are more similar than two arbitrary genotypes in most maps, especially for the maps initialised from the exponential and log-normal distributions.

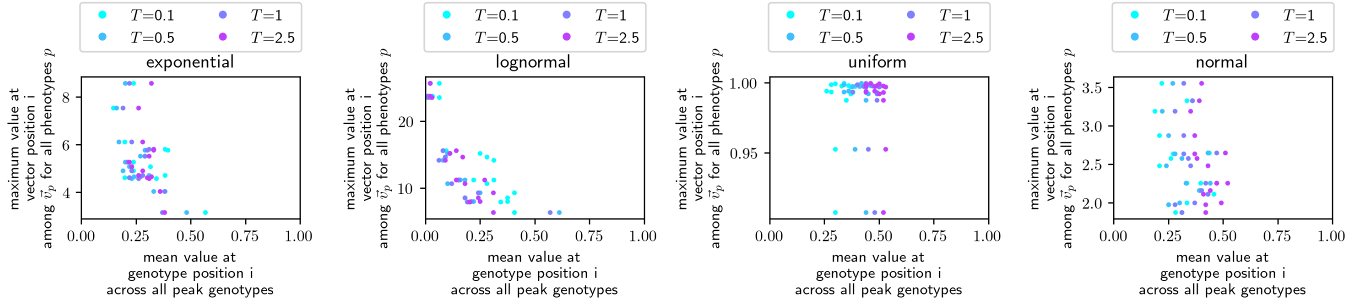

Figure S15: **Why are the 'peaks' for different phenotypes more clustered in genotype space in some ND GP maps (Fig S14)?** Here, the following hypothesis is tested: if one vector element $\vec{v}_p^i$ is very large, then phenotype $p$ has a low value of $G$ and thus a high frequency whenever the $i^{th}$ genotype position is one. Thus, the genotype maximising $P(q|g)$ for any $q \neq p$ is likely to have a zero at the $i^{th}$ genotype position, and the genotypes maximising different $q$ resemble each other more than arbitrary sequences. To test this, the maximum absolute vector element $\vec{v}_r^i$ at position $i$ (across all phenotypes $r$) is plotted against the mean value of all 'peak' genotypes at position $i$. Consistent with the hypothesis, 'peak' genotypes have a compositional bias towards zero, particularly when there is one phenotype $p$ with a particularly high vector element $\vec{v}_r^i$ at position $i$. These high-$\vec{v}_r^i$ values are found especially in the maps initialised from exponential and log-normal distributions.

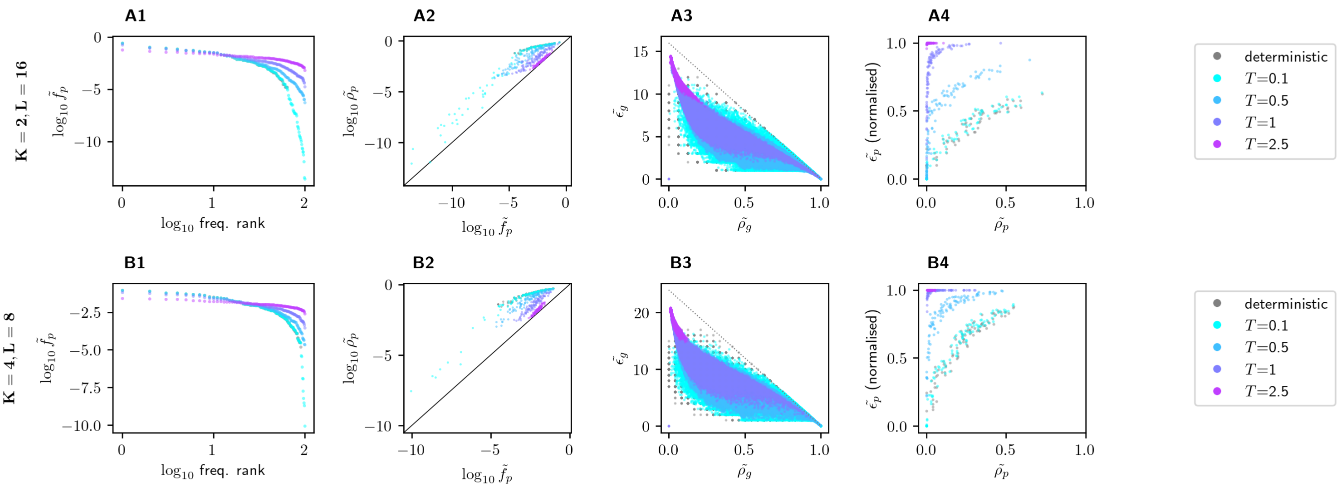

Figure S16: **ND GP map analysis of alternative versions of the synthetic ND GP map, with different alphabet size $K$:** The sequence length $L$ is adjusted to keep the total number of genotypes $K^L$ constant **(row A)** alphabet size $K = 2$ and sequence length $L = 16$, **(row B)** alphabet size $K = 4$ and sequence length $L = 8$. All maps have fixed $n_p = 100$, whereas stochasticity $T$ is varied (with the deterministic limit, where each genotype maps to the lowest-$G$ structure shown in grey for comparison). As in Fig 5 in the main text, each column analyses one specific aspect of the ND GP map: **(1)** The phenotypic frequency $\tilde{f}_p$ is plotted against the frequency rank, showing phenotypic bias. **(2)** The phenotypic robustness $\log \tilde{\rho}_p$ is plotted against the phenotypic frequency $\log \tilde{f}_p$ ($x = y$ is drawn as a black line to guide the eye). Since $\tilde{\rho}_p > \tilde{f}_p$ for the majority of phenotypes, the maps have genetic correlations, but this difference is less pronounced in the high-$T$ versions. **(3)** The genotypic evolvability $\tilde{e}_g$ is plotted against the genotypic robustness $\tilde{\rho}_g$. There are no high-$\tilde{e}_g$-high-$\tilde{\rho}_g$ genotypes, consistent with the upper bound from section S1.2 (grey dotted line; the data is below the bound when accounting for numeric errors of up to $10^{-4}$). **(4)** The phenotypic evolvability $\tilde{e}_p$ is plotted against the phenotypic robustness $\tilde{\rho}_p$. This relationship is generally positive, except for saturation effects, especially in the high-$T$ limit.

#### S5 Threshold-based framework for ND GP maps

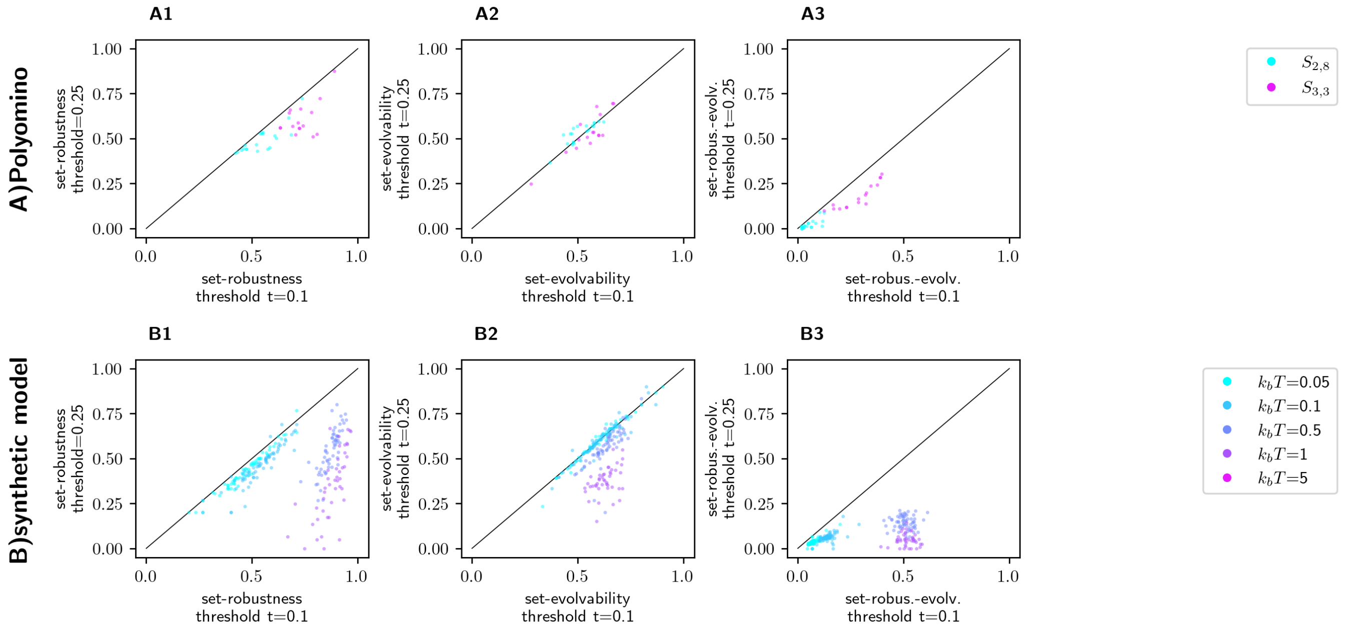

Figure S17: **Analysis of ND GP maps with a threshold-based framework from ref [23]:** the threshold-based framework is applied to the Polyomino map, where it was first proposed **(row A)**, and to the synthetic map with $n_p = 100$ **(row B)**. Since this framework has a free parameter, the threshold $t$ for including a phenotype in the 'set' of a genotype (if $P(p|g) > t$), the sensitivity to this threshold is tested: in **column 1**, the phenotypic set-robustness computed with $t = 0.25$ is plotted against that computed with $t = 0.1$, and similarly for the set-evolvability in **column 2** and the set-robustness-evolvability in **column 3**. The data show that all three measures can be sensitive to the choice of threshold. Thus, this framework is more useful for a specific scenario with known threshold than for charactering a ND GP map in general.

For completeness, let us consider an alternative way of characterising robustness and evolvability in ND GP maps: 'set-robustness' and 'set-evolvability' first proposed by Jouffrey, Leonard & Ahnert [23]. These quantities assume that there is a fixed probability cut-off above which phenotypes are relevant: all phenotypes above the cut-off are treated as equal (even if they have different ensemble frequencies), and all phenotypes below the cut-off are ignored. This was proposed in the context of gene duplication, and it was assumed that correctly folded structures will contribute to fitness if they are more frequent than a given cut-off, but that above this cut-off the exact value of their ensemble frequency is irrelevant. Since it is possible that a mutation both preserves one or more 'relevant' phenotypes, and introduces a new one, this cut-off-based metric can give a positive relationship between evolvability and robustness, even on a genotypic level [23].

Here, the concepts of phenotypic 'set-robustness', 'set-evolvability' and 'set-robust-evolvability' are applied to both the Polyomino ND GP map (where it was first proposed) and the synthetic ND GP map (Fig. S17). Even for a single ND GP map with a fixed set of parameters, 'set-robustness', 'set-evolvability' and 'set-robust-evolvability' are found to be highly dependent on the choice of cut-off in their definitions: when plotting set-robust-evolvability with a cut-off of 0.25 against the set-robust-evolvability of the same phenotype in the same ND GP map but with a cut-off of 0.1 only gives a correlation of $< 0.03$ for the synthetic ND GP maps with $n_p = 100$ and $T \geq 0.5$. Other versions of the synthetic ND GP map (for example, $n_p = 100$ and $T = 0.1$) give much higher correlations. The key factor may be the frequencies found in a typical ensemble: if typical ensemble frequencies are well above/below the cut-off (for example, in low-$T$ ensembles, the most frequent phenotype has an ensemble frequency close to one, and the second-highest a much lower ensemble frequency, see Fig. S12), then the exact value of the cut-off may be less important.

Because of the potential cut-off dependence, these set-based quantities are not used further to characterise ND GP maps. While this paper also uses a cut-off in the Polyomino model for computational reasons (see section S2.5), this cut-off merely discards phenotypes with low ensemble frequencies, but keeps the quantitative ensemble frequency information for all other phenotypes.

#### References

[1] S. F. Greenbury, S. Schaper, S. E. Ahnert, and A. A. Louis, "Genetic correlations greatly increase mutational robustness and can both reduce and enhance evolvability", PLOS Comput. Biol. **12**, e1004773 (2016).

[2] P. García-Galindo, S. E. Ahnert, and N. S. Martin, "The non-deterministic genotype–phenotype map of RNA secondary structure", J. R. Soc. Interface **20**, 20230132 (2023).

[3] A. Sappington and V. Mohanty, "Probabilistic genotype-phenotype maps reveal mutational robustness of RNA folding, spin glasses, and quantum circuits", Phys. Rev. Research **7**, 013118 (2025).

[4] L. W. Ancel and W. Fontana, "Plasticity, evolvability, and modularity in RNA", Journal of Experimental Zoology **288**, 242–283 (2000).

[5] A. Wagner, "Robustness and evolvability: a paradox resolved", Proc. R. Soc. Lond. B **275**, 91–100 (2008).

[6] K. Dingle, S. Schaper, and A. A. Louis, "The structure of the genotype–phenotype map strongly constrains the evolution of non-coding RNA", Interface focus **5**, 20150053 (2015).

[7] J. A. García-Martín, P. Catalán, S. Manrubia, and J. A. Cuesta, "Statistical theory of phenotype abundance distributions: a test through exact enumeration of genotype spaces", EPL **123**, 28001 (2018).

[8] N. S. Martin, C. Q. Camargo, and A. A. Louis, "Bias in the arrival of variation can dominate over natural selection in Richard Dawkins's biomorphs", PLOS Comput. Biol. **20**, e1011893 (2024).

[9] E. Bornberg-Bauer and H. S. Chan, "Modeling evolutionary landscapes: mutational stability, topology, and superfunnels in sequence space", PNAS **96**, 10689–10694 (1999).

[10] E. Bornberg-Bauer, "How are model protein structures distributed in sequence space?", Biophysical journal **73**, 2393–2403 (1997).

[11] S. E. Ahnert, "Structural properties of genotype–phenotype maps", J. R. Soc. Interface **14**, 20170275 (2017).

[12] S. Manrubia, J. A. Cuesta, J. Aguirre, S. E. Ahnert, L. Altenberg, A. V. Cano, et al., "From genotypes to organisms: state-of-the-art and perspectives of a cornerstone in evolutionary dynamics", Physics of Life Reviews **38**, 55–106 (2021).

[13] E. Ferrada and A. Wagner, "A comparison of genotype-phenotype maps for RNA and proteins", Biophys. J. **102**, 1916–1925 (2012).

[14] S. Greenbury and S. E. Ahnert, "The organization of biological sequences into constrained and unconstrained parts determines fundamental properties of genotype–phenotype maps", J. R. Soc. Interface **12**, 20150724 (2015).

[15] M. Weiß and S. E. Ahnert, "Phenotypes can be robust and evolvable if mutations have non-local effects on sequence constraints", J. R. Soc. Interface **15**, 20170618 (2018).

[16] S. Manrubia and J. A. Cuesta, "Distribution of genotype network sizes in sequence-to-structure genotype–phenotype maps", J. R. Soc. Interface **14**, 20160976 (2017).

[17] M. Weiss and S. E. Ahnert, "Neutral components show a hierarchical community structure in the genotype–phenotype map of RNA secondary structure", J. R. Soc. Interface **17**, 20200608 (2020).

[18] M. Weiß and S. E. Ahnert, "Using small samples to estimate neutral component size and robustness in the genotype–phenotype map of RNA secondary structure", J. R. Soc. Interface **17**, 20190784 (2020).

[19] V. O. Pokusaeva, D. R. Usmanova, E. V. Putintseva, L. Espinar, K. S. Sarkisyan, A. S. Mishin, et al., "An experimental assay of the interactions of amino acids from orthologous sequences shaping a complex fitness landscape", PLoS Genet. **15**, e1008079 (2019).

[20] L. Di Bari, M. Bisardi, S. Cotogno, M. Weigt, and F. Zamponi, "Emergent time scales of epistasis in protein evolution", PNAS **121**, e2406807121 (2024).

[21] S. Schaper, I. G. Johnston, and A. A. Louis, "Epistasis can lead to fragmented neutral spaces and contingency in evolution", Proc. R. Soc. Lond. B **279**, 1777–1783 (2012).

[22] A. J. Morrison, D. R. Wonderlick, and M. J. Harms, "Ensemble epistasis: thermodynamic origins of non-additivity between mutations", Genetics **219**, iyab105 (2021).

[23]V. Jouffrey, A. Leonard, and S. Ahnert, "Gene duplication and subsequent diversification strongly affect phenotypic evolvability and robustness", R. Soc. Open Sci. **8**, 201636 (2021).
